# Lowering insulin mitigates female reproductive aging and diet-induced ovarian dysfunction

**DOI:** 10.64898/2025.12.24.696334

**Authors:** Faria Athar, Liam G. Hall, Xiaoke Hu, James D. Johnson, Nicole M. Templeman

## Abstract

Hyperinsulinemia has consequences beyond metabolic dysfunction, including reproductive system effects. We found that hyperinsulinemia at age 46-47 was predictive of earlier menopause in the Study of Women’s Health Across the Nation. To test causality between insulin levels and reproductive aging, we longitudinally evaluated chow- or high-fat, high-sucrose (HFHS)-fed *Ins1*-null female mice with full or partial *Ins2* insulin gene expression. *Ins1^-/-^;Ins2^+/-^*mice had lower HFHS-induced hyperinsulinemia and less weight gain than their full-*Ins2* littermates, despite comparable HFHS-induced glucose intolerance up to 9 months. By 15 months, *Ins1*^-/-^*;Ins2*^+/+^ ovaries showed multinucleated giant cell accumulation with HFHS, while *Ins1^-/-^;Ins2^+/-^* mice were protected against this response and maintained a higher reserve of follicles. Moreover, aged *Ins1^-/-^;Ins2^+/-^* mice were 9-fold more likely to conceive on HFHS than hyperinsulinemic *Ins1*^-/-^*;Ins2*^+/+^ mice. Elevated insulin is therefore a critical mechanistic link between metabolic dysfunction and reproductive aging, and curtailing insulin levels protects against subfertility and HFHS-induced ovarian decline.

## Introduction

The female reproductive system is one of the earliest organ systems to show age-related decline in humans, which has important implications for both fertility and overall health. A finite, non-renewable reserve of 1-2 million oocytes housed in ovarian follicles is present at birth (Faddy and Gosden 1996), and this ovarian reserve gradually depletes over time, culminating in menopause. However, the two decades preceding menopause are associated with reduced fertility, greater risks of pregnancy complications, cycle irregularity, and hormonal dysregulation, demonstrating that reproductive aging begins well in advance of menopause (Davis et al. 2023). In addition to a dwindling ovarian reserve, reproductive aging also entails a decline in oocyte quality, and structural changes such as ovarian fibrosis that contribute towards impairing follicle development and oocyte maturation (Gu et al. 2024; Igarashi et al. 2015). Chronic inflammation and the presence of multinucleated giant cells further disrupt the ovarian immune environment (Converse et al. 2025). Beyond immediate consequences for reproductive health, reproductive aging has significant impacts on cognitive, cardiovascular, immune, bone, and sexual functions. Accordingly, later onset of menopause is associated with longer lifespan and reduced risks of age-related diseases (Hong et al. 2007; Ossewaarde et al. 2005; Shadyab et al. 2017; Xing and Kirby 2024).

While chronological age is a major determinant of female reproductive health, epidemiological and experimental evidence indicates that metabolic dysfunction also influences reproductive outcomes (Athar et al. 2024). For instance, in addition to causing a milieu of metabolic insults, high-caloric high-fat diets are associated with negative repercussions for the female reproductive system (Di Berardino et al. 2024). In humans, excessive calorie intake is linked to ovulatory disorders, infertility, lower fecundity and miscarriage (Bajalan et al. 2019; Chavarro et al. 2007; Garruti et al. 2019; 2019; Gaskins et al. 2019; Grieger et al. 2018; Karayiannis et al. 2018; Willis et al. 2020). Type 2 diabetes is associated with an increased risk of earlier menopause, as are diets high in refined carbohydrates (Dunneram et al. 2018; Khalil et al. 2011; Mehra et al. 2023; Nagel et al. 2005; Sekhar et al. 2015). In mice, high-fat diets lead to decreased litter sizes, anovulation, and increased markers of inflammation in the ovary (Brothers et al. 2010; Hohos and Skaznik-Wikiel 2017; Skaznik-Wikiel et al. 2016; Speakman 2019; Wu et al. 2014). It is challenging to define specific metabolic mechanisms that drive these reproductive system effects, since metabolic disorders or high-fat diets are associated with a broad suite of changes that may include obesity, insulin hypersecretion, insulin resistance, glucose intolerance, and many other shifts in hormone levels and responses. However, insulin is a promising candidate with a key role in the metabolic-reproductive axis.

Elevated insulin, often referred to as hyperinsulinemia, is a cardinal feature of many metabolic disorders that can precede conditions such as obesity, insulin resistance, or hyperglycemia (Carnethon et al. 2002; Ghasemi et al. 2015; Odeleye et al. 1997; Templeman, Skovsø, et al. 2017; Thomas et al. 2019; Tricò et al. 2018; van Vliet et al. 2020). Insulin receptors and other components of the insulin signaling pathway are widely expressed in mammalian ovarian cells, where they modulate follicle survival, selection, and atresia (Castrillon et al. 2003; Acevedo et al. 2007). Insulin receptors are also expressed in GnRH neurons, astrocytes, and Kiss1 neurons, and insulin signalling influences GnRH release and gonadotropin secretion (Brüning et al. 2000; Poretsky et al. 1999; Qiu et al. 2015; 2013). Insulin receptor knockout in pituitary gonadotroph cells or ovarian theca cells partially protects five-month-old mice from high-fat diet-induced infertility (Brothers et al. 2010; Wu et al. 2014). Conversely, experimentally inducing hyperinsulinemia and insulin resistance via exogenous insulin treatment reduces the number and quality of oocytes retrieved after superovulation in young mice (Ou et al., 2012). Given that mean insulin levels have been continually and significantly rising in human populations across the past decades, even among adults without diabetes (Li et al. 2006; Johnson and Churilla 2025), it is important to know how hyperinsulinemia affects the age-related decline in female reproductive function.

In this study, we probed the associations between insulin levels and reproductive aging in clinical data, and tested the causal role of hyperinsulinemia by specifically reducing production of the insulin ligand in female mice. Previous work showed that suppressing high-fat diet-induced hyperinsulinemia in female *Ins1^-/-^;Ins2^+/-^* mice resulted in attenuated obesity, enhanced insulin sensitivity with age, and extended lifespan (Templeman, Flibotte, et al. 2017). Here, we hypothesized that elevated insulin levels mediate harmful effects of high-fat, high-sucrose diets on the female reproductive system, and that hyperinsulinemia causally accelerates reproductive aging.

## Results

### High perimenopausal insulin levels are predictive of an earlier age at menopause

Premenopausal type 2 diabetes has been linked to earlier menopause and primary ovarian insufficiency (Collins et al. 2017; Mehra et al. 2023; Sekhar et al. 2015), and an atypically early onset of menopause has also been associated with concurrent or subsequent differences in glucose homeostasis, insulin levels, and insulin sensitivity (Savukoski et al. 2021; Roa-Díaz et al. 2023). We wished to build on these observations by evaluating whether differences in levels of insulin itself, prior to menopause, are predictive of the subsequent timing of menopause within a wide population of people who are not being treated for diabetes. To test this relationship, we performed secondary analyses of publicly available data from the Study of Women’s Health Across the Nation (SWAN), a community-based study that longitudinally collected biological, behavioral, and socioeconomic information from a diverse population of middle-aged women transitioning through menopause in the United States. We used linear regression models to test whether fasting insulin levels at 46 or 47 years of age (an early data point for this study, which recruited 42-52 year-old participants for the baseline visit) were associated with age at final menstrual period (Table S1). We found that higher perimenopausal fasting insulin was significantly associated with an earlier menopause (β = –0.471, *p* = 0.033; Fig. 1A) after adjusting for metabolic covariates of body mass index (BMI) and fasting glucose levels at the same 46-47 year-old time point, as well as for self-reported race/ethnicity, income strata, and smoking status, demographic covariates with established effects on age of menopause (Schoenaker et al. 2014; Guan et al. 2025). The relationship between insulin and menopause timing may be masked by confounding impacts of obesity, since higher body mass has instead been correlated with a later age of final menstrual period (Gold et al. 2013). However, after accounting for these confounders, the average age of final menstrual period for the hyperinsulinemic group in the highest insulin quartile (≥11.2 μU/mL fasting insulin) was 6-7 months earlier than those in the three lower insulin quartiles (Fig. 1B; Table S2).

**Fig 1.**
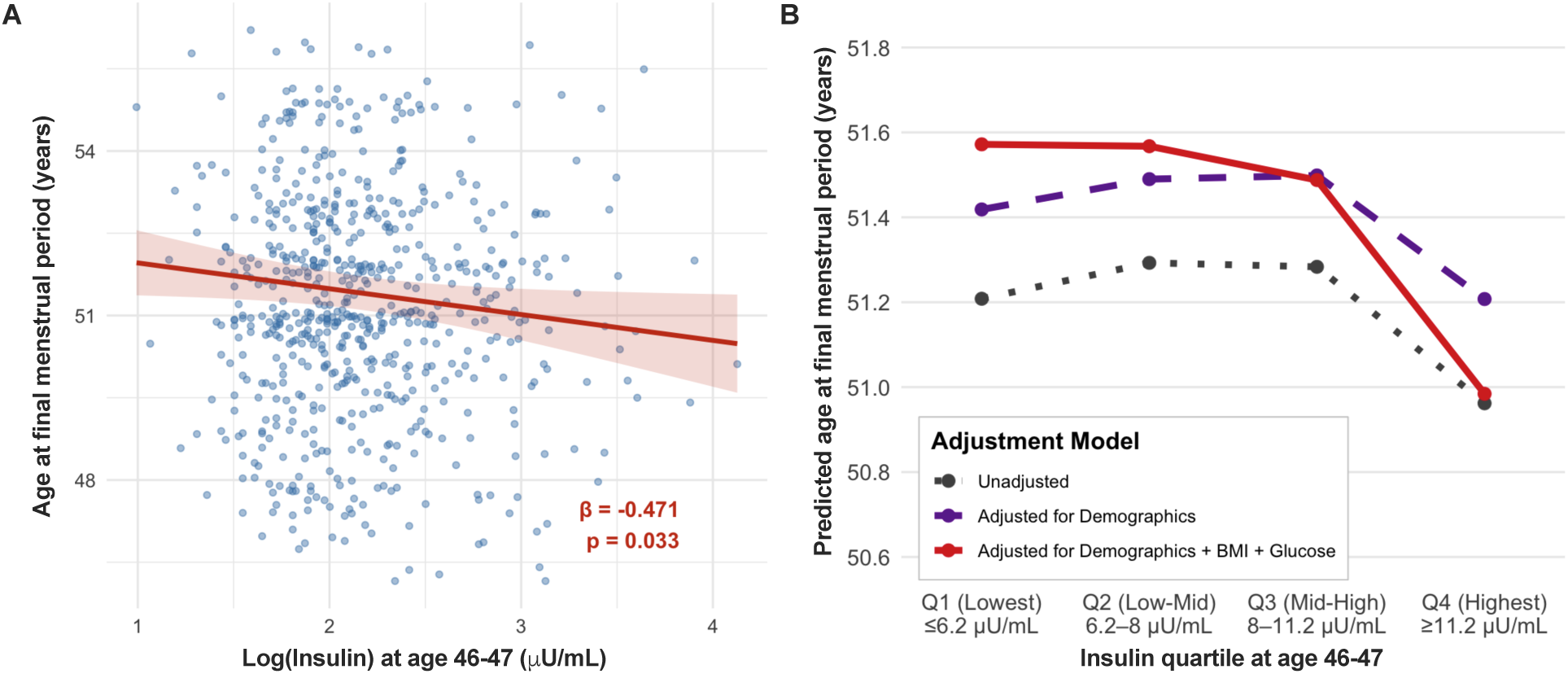
High fasting insulin levels at age 46-47 predict an earlier age at menopause onset. (**A**) Higher levels of log(insulin) at age 46-47 are independently associated with an earlier age of menopause onset (*i.e.*, age of final menstrual period, preceding at least 12 months of amenorrhea), with scatter points showing individual participants in the Study of Women’s Health Across the Nation (SWAN; N = 646). Shaded areas represent 95% confidence intervale around age of final menstrual period in a model fully adjusted for BMI, glucose levels, smoking status, self-reported race/ethnicity, and income. (**B**) The unadjusted relationship, adjustments for demographic variables (self-reported race/ethnicity, smoking status, and income), and additional adjustments for confounding effects of BMI and glucose levels, corresponding to Table S2.

These data revealed an association between elevated insulin levels during perimenopause and an earlier age of natural menopause. However, correlative analyses have limited utility for resolving the nature of this relationship, particularly in light of the myriad other metabolic features that are physiologically interlinked with hyperinsulinemia (Thomas et al. 2019) and may themselves be contributing to menopause timing. Additionally, this midlife dataset provided few insights into the earlier stages of reproductive aging. Therefore, we used mice with genetically reduced insulin to test causality between elevated insulin and accelerated reproductive aging.

### Reduced *Ins2* gene dosage lowers HFHS-induced hyperinsulinemia and obesity, and improves HFHS-induced glucose intolerance with age

To directly test the causal role of hyperinsulinemia in female reproductive aging, we longitudinally evaluated mice with differing levels of insulin production. Specifically, we utilized a mouse model in which littermates had either full or partial expression of the ancestral insulin gene, *Ins2* (*Ins2*^+/+^ versus *Ins2*^+/-^) to modulate endogenous insulin within a normal physiological range of production; all mice had complete inactivation of the rodent-specific *Ins1* gene to prevent compensatory *Ins1* expression. We evaluated mice housed at two different animal facilities to gauge the reproducibility of our *in vivo* observations. Within each facility, female *Ins1^-/-^;Ins2^+/-^* mice and their *Ins1*^-/-^*;Ins2*^+/+^ littermate controls were fed a high-energy, 45%-fat, high-sucrose (HFHS) diet or a 10%-fat, no-sucrose chow diet from weaning onward, and monitored over the subsequent 15 months (Fig. 2A-B). We focused our analyses around 2-3 months (Fig. 2C-F), 9-10 months (Fig. 2G-J), and 14-15 months of age (Fig. 2K-N), as these three time periods approximately represent young adulthood, an early stage of reproductive decline, and a point that is late in the fertility window for inbred mice (Balough et al. 2024; Fig. 2A). Consistent with the genetic perturbation and prior phenotyping (Templeman, Flibotte, et al. 2017; Templeman et al. 2015), *Ins1^-/-^;Ins2^+/-^*mice tended to exhibit slightly lower fasting insulin levels than *Ins1*^-/-^*;Ins2*^+/+^ mice at 2 months of age (Fig. 2C; p = 0.0503 for genotype effect), with a significant reduction evident by 9 months of age across both animal facilities (Fig. 2G; Fig S1). This was particularly pronounced on a hyperinsulinogenic HFHS diet, as HFHS-fed *Ins1^-/-^ ;Ins2^+/-^* mice averaged ∼60% lower fasting insulin levels in circulation than their *Ins1*^-/-^*;Ins2*^+/+^ HFHS-fed counterparts. By 14 months of age, HFHS-fed *Ins1*^-/-^*;Ins2*^+/+^ mice had developed marked fasting and glucose-stimulated hyperinsulinemia (Fig. 2K, 2N), whereas HFHS-fed *Ins1^-/-^;Ins2^+/-^* mice remained relatively protected against diet-induced hyperinsulinemia, with average fasting and glucose-stimulated insulin levels that were reduced by 46% and 37%, respectively.

**Fig 2.**
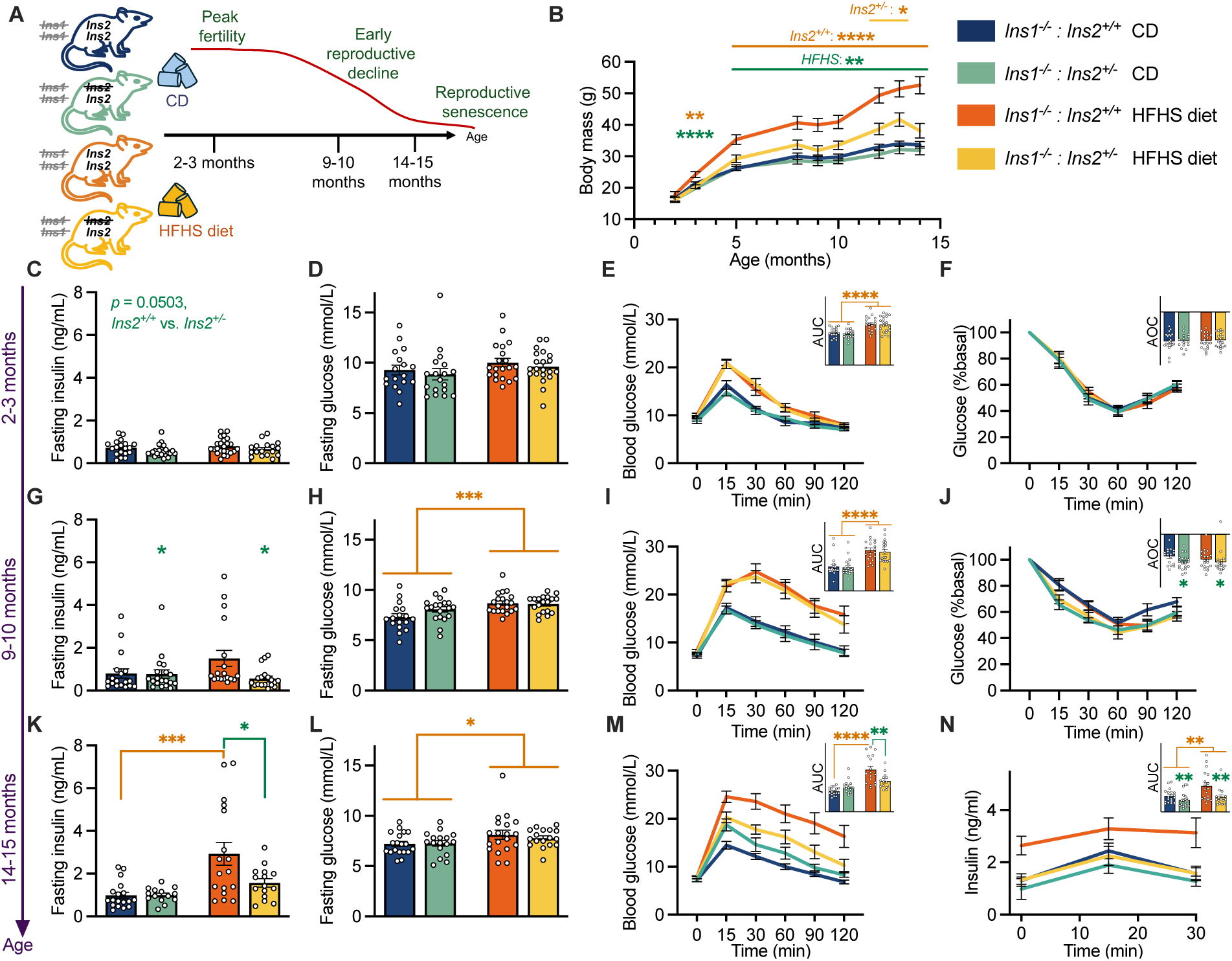
Suppressing hyperinsulinemia reduces weight gain, and eventually improves HFHS diet-induced glucose intolerance in aged female mice. (A) Schematic of the study design. *Ins1*-null mice with either full or partial *Ins2* gene dosage (*Ins2^+/+^* versus *Ins2^+/-^* littermates) were tracked longitudinally up to 15 months, focusing on 2-3 month, 9-10 month and 14-15 month timepoints to assess metabolic and reproductive health. (B) Longitudinal body weight measurements. (C, G, K) Fasting plasma insulin levels at 2-3, 9-10, and 14-15 months of age. (D, H, L) Fasting blood glucose concentrations at 2-3, 9-10, and 14-15 months of age. (E, I, M) Blood glucose response to an intraperitoneal injection of glucose (2 g/kg) at indicated ages. (F, J) Blood glucose response to an intraperitoneal injection of an insulin analog (0.75 U/kg) at indicated ages, and (N) glucose-stimulated insulin secretion measured at 13 months of age. Data are presented as means and SEM; scatter points represent individual mice. Statistical analyses were performed using two-way ANOVA to evaluate the effects of genotype and diet (performed at each age point for body mass data). When significant interactions were detected, one-way ANOVAs with Bonferroni post hoc corrections were conducted. Orange asterisks indicate a statistically significant effect of diet, within one of the genotypes where indicated; green asterisks indicate a statistically significant genotype effect, within one of the diets where indicated. * *p* < 0.05 or *p* < 0.025 with Bonferroni correction (panel K); ** *p* < 0.01; *** *p* < 0.001; **** *p* < 0.0001. n =15-20 mice in total across all groups and experiments (n = 4-11 from Facility A; n = 8-10 from Facility B). Corresponds to Figure S1.

We longitudinally assessed body mass as well as glucose and insulin tolerance to understand the consequences of HFHS feeding and *Ins2* gene dosage in these colonies. Consistent with prior studies (Templeman, Flibotte, et al. 2017; Templeman et al. 2015), hyperinsulinemic *Ins1*^-/-^*;Ins2*^+/+^ mice on the HFHS diet gained significantly more weight than their *Ins1^-/-^;Ins2^+/-^*littermates on the same diet, at both facilities (Fig. 2B, S1A, S1B). Average fasting glucose levels remained comparable between the *Ins2*^+/+^ and *Ins2*^+/-^ genotypes across the duration of this study, with slight HFHS-induced fasting hyperglycemia appearing with age (Fig. 2D, 2H, 2L). 2 month-old HFHS-fed mice showed modest glucose intolerance compared to chow-fed mice, with no difference between genotypes (Fig. 2E). By 9 months of age glucose intolerance was considerable for all HFHS-fed mice, but there was still little difference between genotypes (Fig. 2I). Therefore, we were able to observe effects of specifically reducing insulin—without genotype-associated changes to glucose levels—up to 9 months of age, as *Ins1^-/-^;Ins2^+/-^* mice did not overtly differ in their degree of HFHS-induced hyperglycemia or glucose intolerance to this point. However, some enhancement in insulin sensitivity was becoming evident in *Ins1^-/-^;Ins2^+/-^*mice at 9 months of age (Fig. 2J), consistent with prior observations that hyperinsulinemia helps perpetuate insulin resistance (Templeman, Flibotte, et al. 2017). Limiting insulin hypersecretion and the consequential improvements for phenotypes such as insulin sensitivity eventually extended beneficial effects on glucose homeostasis in these mice, as 14 month-old *Ins1^-/-^;Ins2^+/-^*mice on the HFHS diet had superior glucose tolerance compared to hyperinsulinemic *Ins1*^-/-^*;Ins2*^+/+^ mice on the same diet (Fig. 2M). Overall, our metabolic data across both facilities indicate that compared to their full-*Ins2* littermates, *Ins1^-/-^;Ins2^+/-^*mice had lower insulin levels and less HFHS-induced weight gain, and they exhibited delayed improvements to HFHS-induced glucose intolerance by the age of 14 months, despite similar degrees of glycemia and HFHS-associated glucose intolerance up to 9 months.

### Lowering insulin slows the depletion of the ovarian reserve and protects against multinucleated giant cell accumulation in aged ovaries

To characterize impact of insulin levels on ageing of the female reproductive system, we examined murine ovarian reserve and other age-related features of ovarian architecture. Follicle number and circulating anti-Müllerian hormone (AMH) levels are both established markers of ovarian reserve that decline with advanced age in mice and humans (Hall 2015; Balough et al. 2024). While AMH levels showed considerable variability within each group, the HFHS-fed *Ins1*^-/-^*;Ins2*^+/+^ mice tended to have the lowest average AMH levels compared to other experimental groups at 9-10 and 14-15 months of age, although this trend was not statistically significant (Fig. 3B, 3I). Consistent with this pattern, total follicular reserve was significantly reduced in *Ins1*^-/-^*;Ins2*^+/+^ ovaries compared to *Ins1^-/-^;Ins2^+/-^* ovaries at 9 and 15 months of age (Fig. 3C, 3J). After staging the ovarian follicles, it seemed that the differences between genotypes were most pronounced for secondary follicles at 9 months (Fig. 3F) and for primary follicles at 15 months (Fig. 3L), both of which were significantly reduced in *Ins1*^-/-^*;Ins2*^+/+^ ovaries compared to *Ins1^-/-^;Ins2^+/-^* ovaries. Because each ovulated oocyte typically gives rise to a single corpus luteum (CL), we quantified CL numbers as an additional metric of ovarian functioning. Unlike total follicle counts, corpora lutea at 9 months were significantly influenced by diet, signifying that HFHS-fed mice likely underwent fewer ovulations (Fig. 3D). Together, these data suggest that insulin levels affect ovarian follicle depletion, and lowering *Ins2* gene dosage in female mice preserves the ovarian reserve.

**Fig 3.**
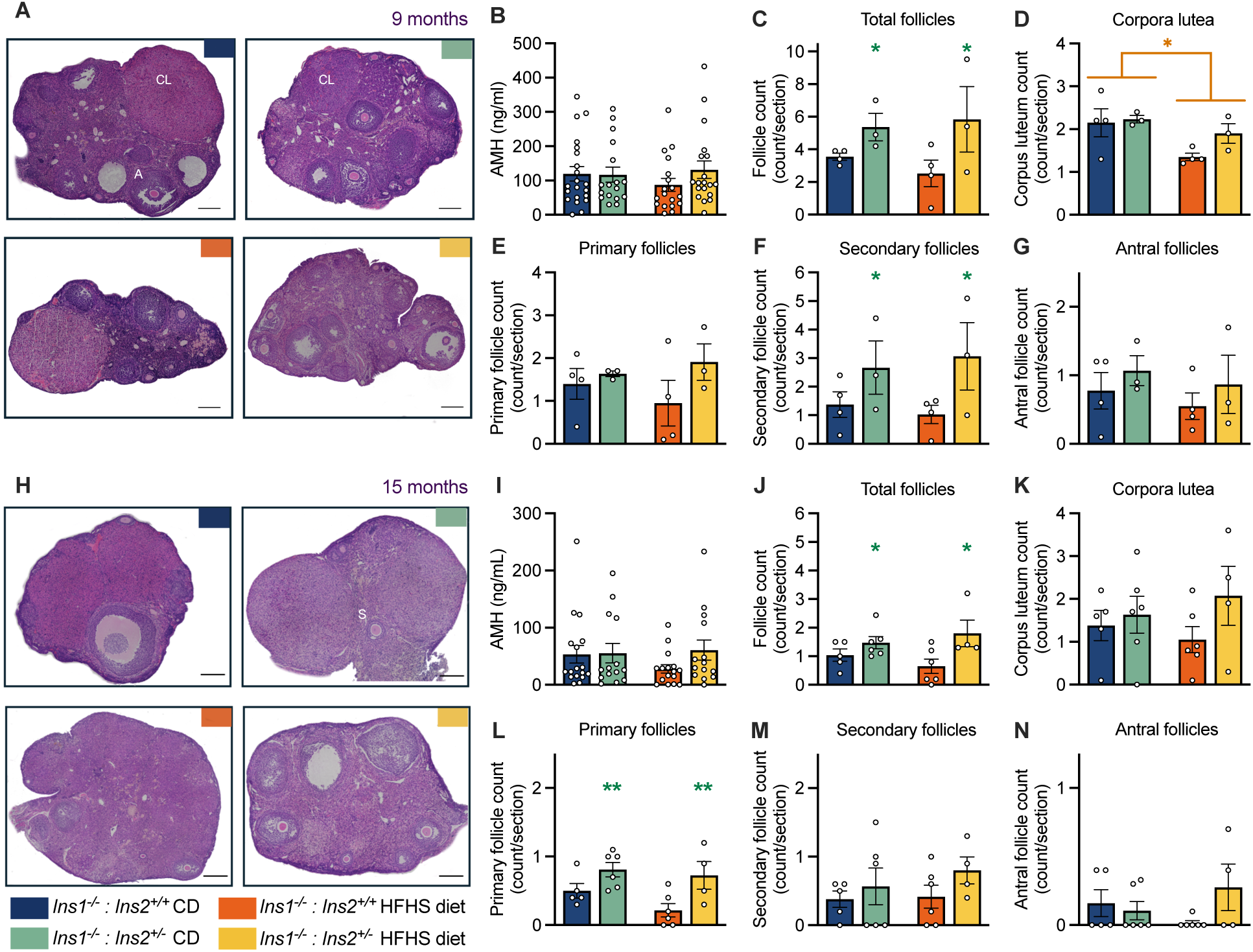
*Ins1*^-/-^*;Ins2*^+/+^ mice have fewer ovarian follicles at 9 and 15 months of age. (A) Representative images of H&E-stained ovarian sections from 9-month-old mice. CL = corpora lutea; A = antral follicle. (B–G): Plasma AMH levels (B), total follicle counts (C), corpora lutea (D), primary follicles (E), secondary follicles (F), and antral follicles (G) in 9-month-old mice. n = 17-20 mice for AMH and n = 3-4 mice for ovarian histology. (H) Representative images of H&E-stained ovarian sections. S = secondary follicle. (I–N): Plasma AMH levels (I), total follicle counts (J), corpora lutea (K), primary follicles (L), secondary follicles (M), and antral follicles (N) in 15-month-old mice. n = 14-17 mice for AMH and n = 4-6 mice for ovarian histology. All mice for these assessments were housed in Facility A. Data are presented as means and SEM; scatter points represent individual mice. Two-way ANOVAs were performed to evaluate the effects of genotype and diet. Orange asterisks indicate a statistically significant effect of diet; green asterisks indicate a statistically significant genotype effect. * *p* < 0.05; ** *p* < 0.01. Scale bars = 400 μm.

Aging ovaries exhibit characteristic structural changes, including hemorrhagic corpora lutea, multinucleated epithelial giant cell accumulation, and fibrosis. To evaluate whether insulin levels affect these aging features, we conducted further gross histological analyses of ovaries from 15 month-old mice and observed structural abnormalities that appeared linked to HFHS-induced hyperinsulinemia. For example, we noted that a subset of HFHS-fed *Ins1*^-/-^*;Ins2*^+/+^ mice tended to have a relatively higher proportion of corpora lutea with haemorrhage, compared to other groups (Fig. 4A-B), although hemorrhagic corpora lutea were generally a rare observation. Hemorrhagic corpora lutea are a marker of defective ovulatory wound healing (Cheng et al. 2020; Zaniker et al. 2023; Duffy et al. 2019), so chronic hyperinsulinemia may contribute to specific features of ovarian aging. We also assessed the prevalence of multinucleated epithelial giant cells (MNGCs), which are a unique immune cell population formed by macrophage fusion that are associated with ovary tissue degradation and immune signalling (Converse et al. 2025; Foley et al. 2021). MNGCs are a sign of inflammaging, and their presence in aged ovaries signifies the likelihood of poor debris-clearing mechanisms (Converse et al. 2025; Balough et al. 2024). The HFHS diet markedly increased MNGC prevalence in aged ovaries of *Ins1*^-/-^*;Ins2*^+/+^ mice (p = 0.0128), whereas the ovaries of HFHS-fed *Ins1^-/-^;Ins2^+/-^*mice had significantly fewer MNGCs than their hyperinsulinemic littermates (p = 0.018) and did not show a HFHS-induced increase in MNGC presence (Fig. 4C-D). Fibrosis is a marker of ovarian aging that stiffens tissues and compromises ovulation (Umehara et al. 2022). We did not detect statistically significant differences in levels of ovarian fibrosis across groups at either 9 months or 15 months of age (Fig. 4E-H), but marked age-related increases in ovarian fibrosis may not develop until around 18 months in mice (Briley et al. 2016). All together, our observations of reduced follicle numbers, hemorrhagic CL morphology, and multinucleated epithelial giant cell accumulation suggest that hyperinsulinemia accelerates multiple features of ovarian aging in HFHS-fed *Ins1*^-/-^*;Ins2*^+/+^ mice.

**Fig 4.**
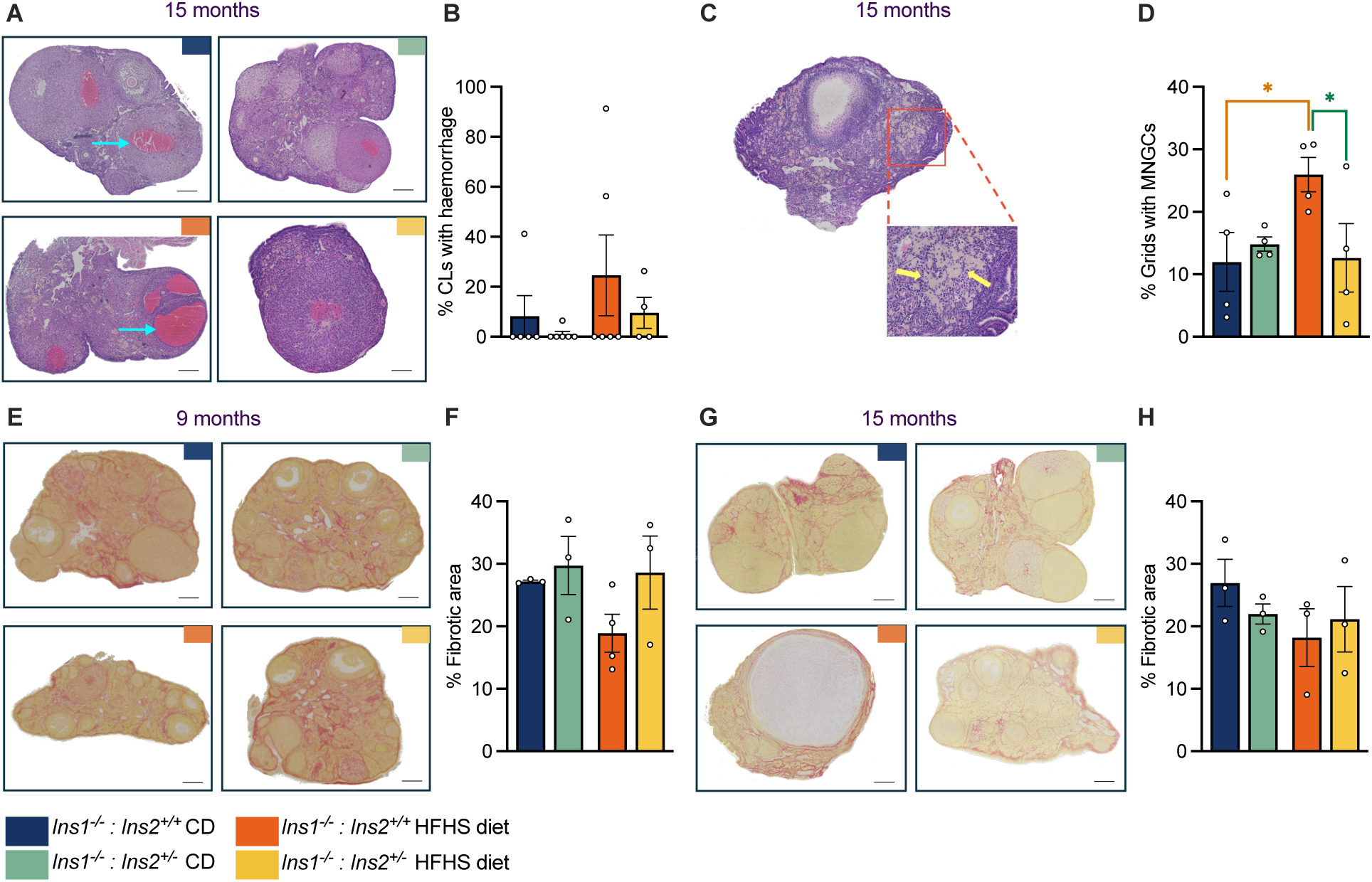
Hyperinsulinemia alters ovarian architecture features in reproductively-aged mice. (A) Representative H&E-stained ovaries from 15-month-old mice showing signs of haemorrhaging within the ovary, with blue arrows indicate haemorrhaging, and (B) quantification of the percent of CLs that were haemorrhagic per mouse. n = 4–6 mice per group. (C) Representative H&E sections with stromal regions of the ovary magnified. Yellow arrows point toward foamy, brownish MNGCs. (D) Quantification of multinucleated epithelial giant cell (MNGC) prevalence in grids overlayed on ovary images that were blinded and randomized. n = 4 mice per group. (E–F) Representative PSR-stained images of ovaries from 9-month-old mice and quantification of PSR-positive area (n = 3–4 mice per group). (G–H) Representative PSR-stained sections from 15-month-old mice and quantification of PSR-positive area (n = 3–4 mice per group). All mice for these assessments were housed in Facility A. Data are presented as means and SEM; scatter points represent individual mice. Orange asterisks indicate a statistically significant effect of diet; green asterisks indicate a statistically significant genotype effect. * *p* < 0.025. Scale bars = 400 μm.

### Reductions in *Ins2* and circulating insulin did not lead to lasting estrous cycle changes

Circulating FSH levels rise during aging as a compensatory response to declining ovarian function and reduced negative feedback from the ovaries, while estrous cycles become irregular with age (Hall 2015). Insulin receptor signaling in hypothalamic GnRH neurons and/or pituitary gonadotroph cells is linked to central regulation of the hypothalamus-pituitary-gonad axis, including effects on GnRH pulsatility, gonadotropin levels, and estrous cyclicity (DiVall et al. 2015; Brothers et al. 2010). Therefore, we examined estrous cycle patterns and FSH levels periodically across the study time course. We observed that 2-3 month-old *Ins1*^-/-^*;Ins2*^+/+^ mice spent more time in diestrus and less time in metestrus compared to *Ins1^-/-^;Ins2^+/-^*mice, across both diets, but these modest differences were no longer evident at later age points (Fig. 5A, 5C, 5E). There were also no strong effects of the 45%-fat, high-sucrose diet on estrous cycling.

**Fig 5.**
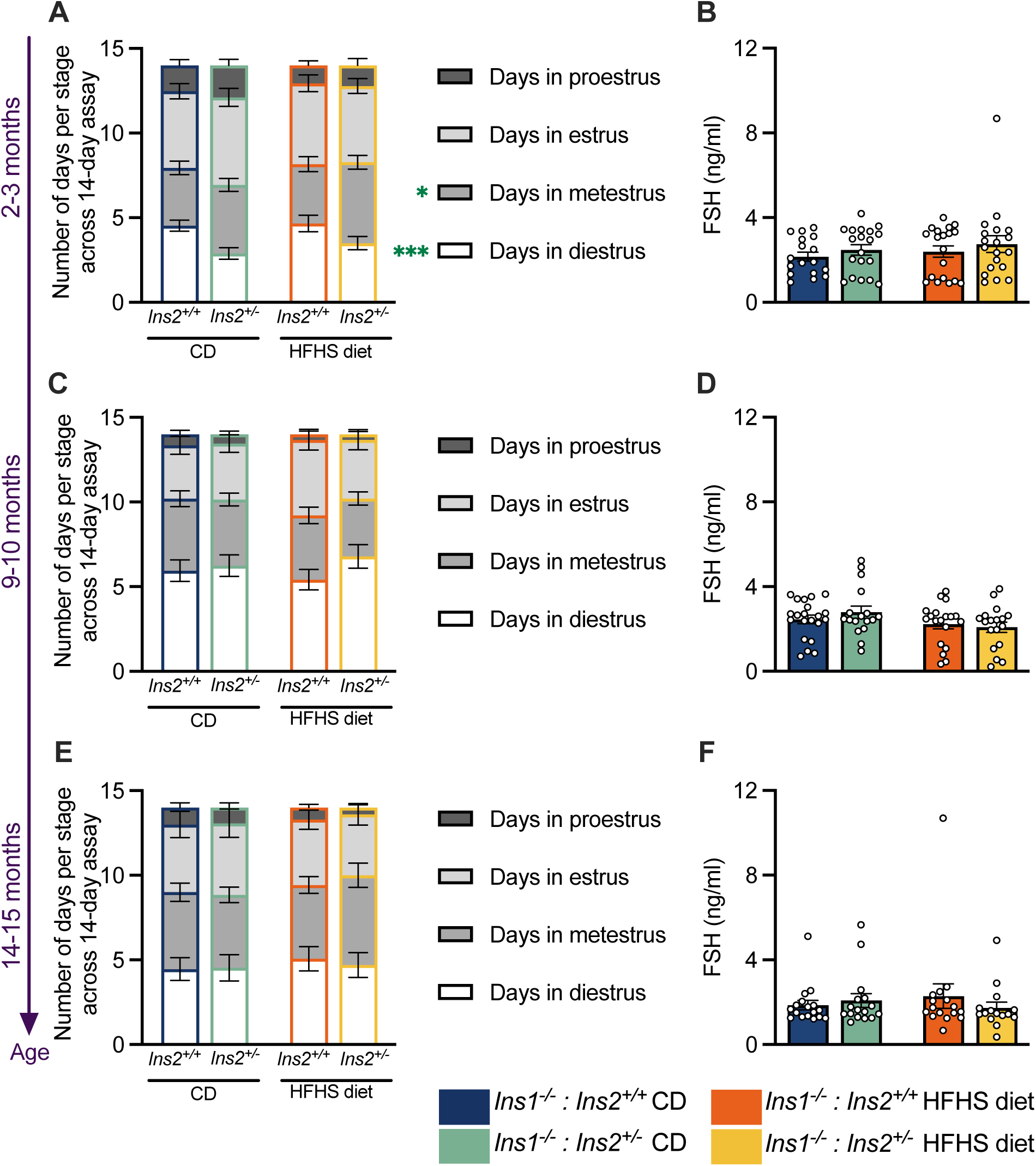
Estrous cycles and FSH remain consistent across experimental groups. Number of times mice cycled through each stage of the estrous within a 14-day period, at (A) 2-3 months of age, (C) 9-10 months of age, and (E) 14-15 months of age. (B, D, F) Plasma FSH levels in female mice at 2-3, 9-10 and 14-15 months of age, respectively, n = 15-20 mice. Data are presented as means and SEM; scatter points represent individual mice. Statistical analyses were performed using two-way ANOVAs to evaluate the effects of genotype and diet. Green asterisks indicate a statistically significant genotype effect. * *p* < 0.05; *** *p* < 0.001.

These observations indicate that estrous cycle disruptions were unlikely to be a major contributor to diet- or insulin-induced differences in age-related reproductive decline in these mice. Plasma FSH levels typically begin to increase around 12-16 months of age across all estrous stages in mice, even when the ovarian reserve of primordial follicles remains substantial (Balough et al. 2024; Parkening et al. 1982). However, we observed no significant differences between *Ins1*^-/-^*;Ins2*^+/+^ and *Ins1^-/-^;Ins2^+/-^*mice in FSH levels during diestrus at any of the three time points tested, nor any pronounced diet effects (Fig. 5B, 5D, 5F). Together, these data suggest that *Ins2* gene dosage and circulating insulin levels may directly affect ovaries to a larger extent than central components of the hypothalamus-pituitary-gonad axis during reproductive aging.

### Lowering endogenous insulin levels preserves fertility in aging female mice

A major goal of our study was to test whether curtailing insulin production could be sufficient to slow the age-related decline in fertility. To address this, we longitudinally assessed fertility and pregnancy outcomes by periodically pairing the female mice with young (2-month old) wild-type C57BL/6 males. At the young, 2-3-month timepoint, all female mice became successfully pregnant (Fig. 6A) and there were comparable full-term delivery rates across all groups (Fig. 6B). This indicates robust early reproductive function across both genotypes, regardless of diet, suggesting that *Ins2* gene dosage did not significantly alter baseline reproductive parameters. Young HFHS-fed mice averaged fewer pups per litter than CD-fed mice (Fig. 6D), but litter size did not appear to contribute to diet-induced fertility differences for 9-10-month-old mice (Fig. 6H). By 9-10 months of age, *Ins1*^-/-^*;Ins2*^+/+^ female mice with higher insulin levels tended to experience pregnancy loss more frequently than *Ins1^-/-^;Ins2^+/-^* mice. Across both facilities combined, the live birth rate of pregnant *Ins1^-/-^;Ins2^+/-^* mice was approximately two times higher than the live birth rate of pregnant *Ins1*^-/-^*;Ins2*^+/+^ mice within each diet treatment, and in facility A, *Ins1^-/-^;Ins2^+/-^* mice were the only ones to carry litters to term at the 9-10-month time point (p = 0.0256 on chow diet in Facility A; Fig. 6F). By the 14-15-month timepoint, fertility decline was especially apparent in hyperinsulinemic *Ins1*^-/-^*;Ins2*^+/+^ female mice on the HFHS diet. The average pregnancy rate of HFHS-fed *Ins1*^-/-^*;Ins2*^+/+^ mice was only ∼7% at this reproductively aged time point, significantly lower than both their chow-fed controls (p = 0.0008) and their diet-matched reduced-*Ins2* littermates (p = 0.0044). In contrast, HFHS-fed *Ins1^-/-^;Ins2^+/-^* mice maintained ∼64% pregnancy rates, comparable to the chow-fed group (p = 0.9 for HFHS-fed *Ins1^-/-^;Ins2^+/-^* versus CD-fed *Ins1^-/-^;Ins2^+/-^*; Fig. 6I). Therefore, in consistent patterns across two distinct animal facilities, HFHS-fed *Ins1^-/-^;Ins2^+/-^* mice were approximately 9 times more likely to conceive than their hyperinsulinemic *Ins1*^-/-^*;Ins2*^+/+^ HFHS-fed counterparts. However, of those female mice that conceived, unsuccessful pregnancy outcomes were equally prevalent among 14-15 month-old HFHS-fed mice regardless of genotype (Fig. 6J). Time to birth from pairing remained consistent across all groups at all three ages (Fig. 6C, 6G, 6K).

**Fig 6.**
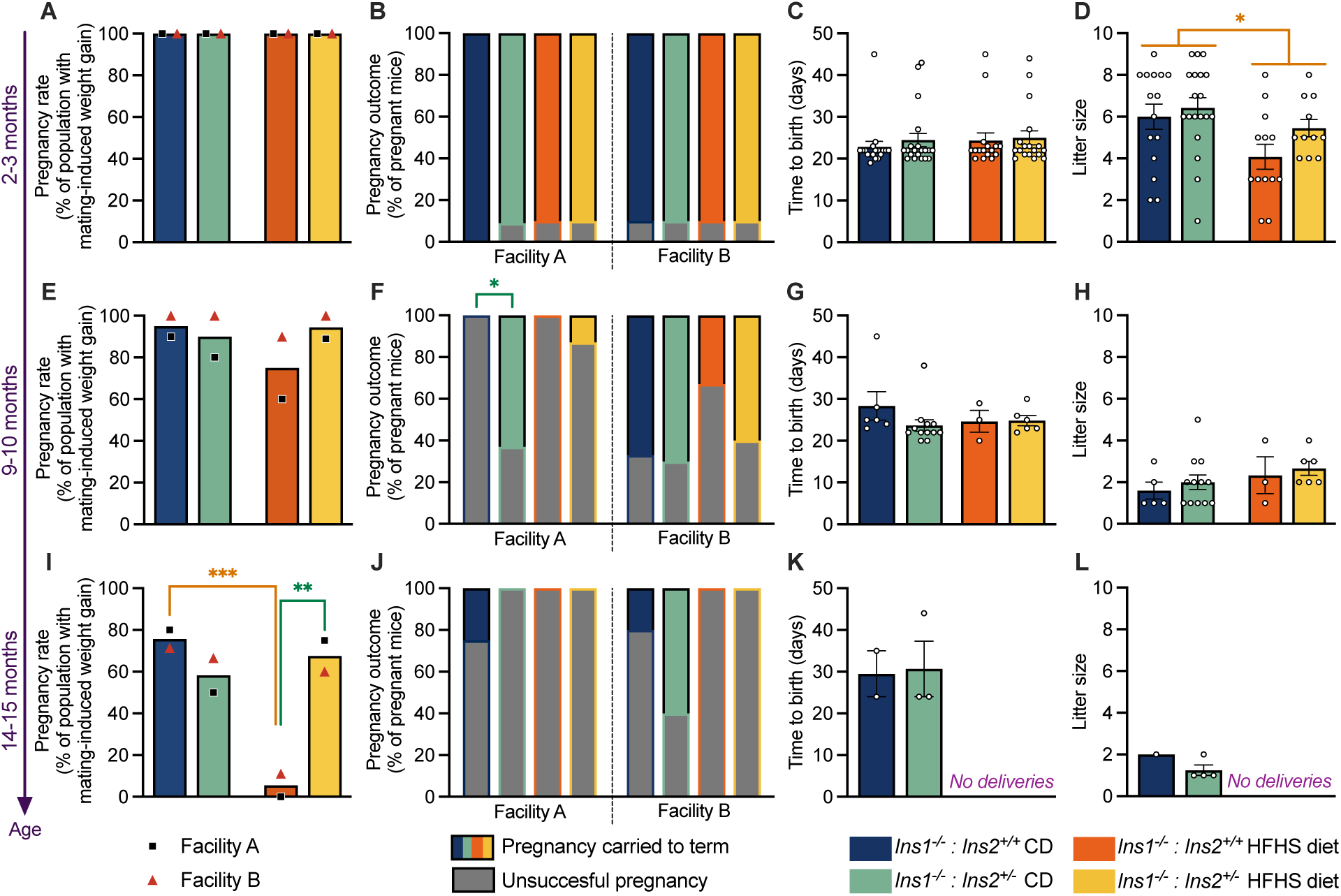
Hyperinsulinemia drives diet-induced infertility in reproductively aged female mice. (A–D) Pregnancy rates, pregnancy outcomes, time to delivery, and litter sizes from breeding trials of female mice at 2-3 months of age (n = 20), (E–H) 9-10 months of age (n = 15-20), and (I–L) 14-15 months of age (n = 12-20). Pregnancy success was monitored during a 4-week mating period with a wild-type 2-month-old male mouse. Data are presented as means and SEM. Scatter points represent individual dams for time to delivery and litter size data (C-D, G-H, K-L). Black squares indicate the average pregnancy rates from facility A, and red triangles indicate average pregnancy rates from facility B (A, E, I). Fisher’s exact test was used to evaluate pregnancy rates and outcomes across groups; two-way ANOVAs were used for time-to-birth and litter size data to evaluate the effects of genotype and diet. Orange asterisks indicate a statistically significant effect of diet; green asterisks indicate a statistically significant genotype effect. * *p* < 0.05; ** *p* < 0.01; *** *p* < 0.001.

Therefore, we observed that an age-related decline in fertility was most pronounced in HFHS-fed *Ins1*^-/-^*;Ins2*^+/+^ mice, likely underpinned by an assortment of ovary-level changes. Conversely, genetically reducing insulin levels appears to preserve ovarian reserve and protect reproductively aged *Ins1^-/-^;Ins2^+/-^* mice against aspects of HFHS-exacerbated subfertility, despite their chronic HFHS feeding. This supports the concept that hyperinsulinemia is a modifiable risk factor for female reproductive aging.

## Discussion

Although it is difficult to dissect the precise effects of elevated insulin from interrelated metabolic features, many links exist between hyperinsulinemia and female reproductive dysfunction. For instance, diets that stimulate a rise in insulin levels, obesity, and metabolic syndrome traits can additionally lead to adverse impacts on fertility, fecundity, and pregnancy outcomes (Bajalan et al. 2019; Chavarro et al. 2007; Garruti et al. 2019; 2019; Gaskins et al. 2019; Grieger et al. 2018; Karayiannis et al. 2018; Willis et al. 2020; Hohos and Skaznik-Wikiel 2017; Skaznik-Wikiel et al. 2016; Speakman 2019). Elevated insulin is also a cardinal feature of pathological female reproductive system disorders such as polycystic ovary syndrome and reproductive system cancers (Cust et al. 2007; Gunter et al. 2008; Sun et al. 2016). Considering that altered metabolic health is associated with changes to menopause timing (Mehra et al. 2023; Wellons et al. 2017; Sekhar et al. 2015), we chose to explore whether hyperinsulinemia itself—which can lie mechanistically upstream of obesity and other components of metabolic dysfunction—shapes the progression of female reproductive aging.

We show that hyperinsulinemia accelerates reproductive aging through mechanisms that appear independent of glucose homeostasis and obesity. Using clinical data from the Study of Women’s Health Across the Nation, we found that on average, women with the highest fasting insulin levels at perimenopause (≥11.2 μU/mL) experienced their final menstrual period approximately 6-7 months earlier than those with lower insulin levels, after accounting for BMI, fasting glucose, and demographic confounders (β = –0.471, *p* = 0.033). While prior studies showed that type 2 diabetes is associated with earlier natural menopause and primary ovarian insufficiency (Collins et al. 2017; Mehra et al. 2023; Sekhar et al. 2015), our findings strengthen this link by isolating fasting insulin as a predictor. Since mean insulin levels have been rising in recent decades, these observations may have significant implications (Li et al. 2006; Johnson and Churilla 2025).

To test the causal relationship between insulin and reproductive aging, we used a mouse model to limit insulin production by decreasing *Ins2* gene dosage on an *Ins1*-null background. We found that a hyperinsulinogenic high-fat, high-sucrose diet accelerates the age-related progression towards infertility and promotes detrimental ovarian aging features. As previously reported (Templeman et al. 2015), female *Ins1^-/-^;Ins2^+/-^*mice in our study had attenuated diet-induced hyperinsulinemia and obesity. We observed that lowering insulin through partial *Ins2* deletion also delayed the age-related loss of fertility in HFHS-fed female mice, as *Ins1^-/-^;Ins2^+/-^*mice were approximately 9 times more likely to conceive than hyperinsulinemic littermates at an advanced reproductive age. Suppressing insulin levels preserved ovarian reserve, evidenced by higher ovarian follicle counts in *Ins1^-/-^;Ins2^+/-^*mice compared to *Ins1*^-/-^*;Ins2*^+/+^ littermates. Additionally, we observed that preventing HFHS-induced hyperinsulinemia lowered the incidence of ovarian aging traits, specifically hemorrhagic corpora lutea and MNGC penetrance.

Our findings extend prior work that implicated a role for insulin in regulating female reproductive function. High-fat, high-sugar diets that elevate insulin signaling and other features of metabolic dysfunction are associated with accelerated ovarian follicle loss and subfertility, even without significant weight differences from controls (Hohos et al. 2018; Hohos and Skaznik-Wikiel 2017; Parkening et al. 1982; Skaznik-Wikiel et al. 2016). This suggests that diet-induced metabolic features (such as hyperinsulinemia) may drive reproductive decline to a greater extent than adiposity per se. Indeed, insulin receptors are expressed in oocytes and on somatic cells of the ovary, where insulin signaling affects steroidogenesis and follicular development (Diamanti-Kandarakis and Dunaif 2012; Poretsky et al. 1999). In *Caenorhabditis elegans* nematodes, genetically reducing insulin-like signaling components significantly delays reproductive aging (Luo et al. 2010; Tissenbaum and Ruvkun 1998). Caloric restriction lowers insulin levels in rodents, among other metabolic effects, and this intervention also mitigates age-related chromosomal and mitochondrial abnormalities in oocytes, maintains ovarian reserve, and improves offspring survival (Garcia et al. 2019; Mishina et al. 2021; Selesniemi et al. 2008). Insulin receptor knockout in ovarian theca cells, GnRH neurons, or the pituitary protect young female mice against the pronounced reproductive dysfunction induced by a 60%-fat diet, indicating that insulin may act through multiple tissues to regulate reproductive health (Wu et al. 2014; DiVall et al. 2015; Brothers et al. 2010). However, interpreting data from insulin receptor knockout models is complicated by the functional overlap between insulin and IGF-1 signaling, as these ligands can activate one another’s receptors and insulin receptors can form hybrid receptors with IGF-1 receptors (Blüher et al., 2002; Kitamura et al., 2003; Nakae et al., 2001; Turvey et al., 2022, Vella et al., 2018). Our approach of using genetically reduced endogenous insulin levels allowed for an unambiguous assessment of how the insulin ligand itself affects reproductive aging.

Due to the role of hyperinsulinemia in promoting obesity (Mittendorfer et al. 2024; Templeman, Skovsø, et al. 2017), the *Ins1^-/-^;Ins2^+/-^*mice in our study were partially protected from HFHS diet-induced weight gain compared to their hyperinsulinemic *Ins1*^-/-^*;Ins2*^+/+^ littermates. This means that our mouse model could not separate the direct effects of hyperinsulinemia on reproductive system phenotypes from the possibility of indirect, obesity-mediated effects. Moreover, despite similar degrees of fasting glycemia and diet-induced glucose intolerance up to 9 months of age, reproductively aged HFHS-fed *Ins1^-/-^;Ins2^+/-^* mice showed improvements in glucose tolerance compared to *Ins1*^-/-^*;Ins2*^+/+^ littermates by 14 months of age. While protections against HFHS-induced metabolic dysfunction were incomplete and delayed in *Ins1^-/-^;Ins2^+/-^*mice, this leaves open the possibility that hyperinsulinemia lowers pregnancy rates through some combination of immediate reproductive system effects and secondary effects of these metabolic responses. Other metabolic hormones or inflammatory mediators that covary with insulin may also be contributing to the reproductive aging phenotypes we observed. Nevertheless, since endogenous insulin production was genetically modulated in our mouse model, our data demonstrate that insulin levels themselves can ultimately influence reproductive aging.

Overall, this study identifies hyperinsulinemia as a key feature of metabolic dysfunction that worsens the age-related deterioration of the female reproductive system. By integrating human epidemiological evidence with controlled rodent experiments, we established that elevated insulin levels causally contribute to earlier reproductive decline and ovarian aging. Conversely, suppressing hyperinsulinemia can effectively slow this decline in reproductive function, sustain fertility with age, and alleviate reproductive system responses to chronic high-fat, high-sucrose feeding. Given that insulin reduction can be achieved through targeted dietary and lifestyle interventions (Ritenbaugh et al. 2003; Bird and Hawley 2017; Kirwan et al. 2009; Brown et al. 2020; Kim and Park 2013), these findings could inform strategies for improving female reproductive longevity and overall health.

## Methods

### Data from the Study of Women’s Health Across the Nation (SWAN)

We performed secondary analyses using publicly available data from The Study of Women’s Health Across the Nation (SWAN), a longitudinal study of midlife women across seven sites in the United States (Sowers et al. 2000). Data were retrieved from the Inter-University Consortium for Political and Social Research/ICPSR archive (ICPSR 28762, ICPSR 29221, ICPSR 29401, ICPSR 29701, ICPSR 30142, ICPSR 30181, ICPSR 30501, ICPSR 31181, ICPSR 31901, ICPSR 32122, ICPSR 32721, ICPSR 32961). The SWAN study protocol was approved by the institutional board at each study site, and all participants provided written informed consent. Our secondary data analyses were approved by the Human Research Ethics Board at the University of Victoria (protocol #25-0295).

The 3302 participants enrolled in SWAN at baseline had an intact uterus and at least one ovary, were not taking hormone therapy, pregnant, or lactating, and had experienced one or more menses in the prior 3 months. We restricted our analyses to participants who attended study visits at the ages of 46 and/or 47 and had complete data for insulin measurements at a minimum of one of those visits. Participants were excluded from our analyses if their survey responses indicated that they were taking insulin therapy or medications to manage blood sugar (n = 357), or if they underwent an oophorectomy or hysterectomy during the course of the study (n = 252), or if they experienced menopause before the age of 46 (n= 30). Age of final menstrual period was calculated from the age of participants’ last self-reported menstrual cycle that preceded at least 12 months of amenorrhea.

Measurements taken in the SWAN protocol have been described in detail (Sowers et al. 2000; Matthews et al. 2009; Mitro et al. 2024). In brief, blood was collected after an overnight fast and centrifuged to collect serum for analysis. Insulin was measured by a solid phase radioimmunoassay using Diagnostic Products Corporation reagents (DPC Coat-a-Count; Diagnostic Products Corporation, Los Angeles, CA, USA). Fasting insulin values were used from study participants when they were 46 years old (n = 37 of analyzed participants) or 47 years old (n = 609 of analyzed participants), depending on data availability. Fasting insulin values were log-transformed to normalize distribution in our analyses. A single insulin measurement per participant was used in our analyses, corresponding with a single fasting glucose value and single BMI measurement from the same age point. Fasting glucose was measured on a Hitachi 747-200 (Roche Molecular Biochemicals Diagnostics, Indianapolis, IN) using a hexokinase-coupled reaction. Body mass index (BMI) was measured at clinic visits using a stadiometer and balance beam, and was determined as weight in kg divided by height^2^ in meters.

### Experimental animals

All animal procedures were approved by the University of Victoria Animal Care Committee (AUP #2021-017) and the University of British Columbia Animal Care Committee (AUP #A21-0135) and conducted in accordance with the Canadian Council for Animal Care guidelines. Our experiments were conducted in two separate cohorts of animals, each cohort housed in one of two institutions: the University of British Columbia (facility A) or the University of Victoria (facility B). Female littermate *Ins1*-null mice with either full (*Ins1*^-/-^*;Ins2*^+/+^) or partial (*Ins1^-/-^ ;Ins2^+/-^*) *Ins2* gene dosage were used. This colony was on a predominantly (>90%) C57BL/6J background (strain generously provided by J. Johnson, original mutations created by J. Jami and colleagues;(Duvillié et al. 1997)). Murine *Ins2* is the ancestral insulin gene that is homologous to human *INS* and contributes about 2/3rds of total insulin production (Mehran et al. 2012; Duvillié et al. 1997; Leroux et al. 2001). Thus, modulating *Ins2* gene dosage through full or partial expression of *Ins2* enabled us to study the specific role of insulin hypersecretion in a mammalian model of reproductive aging. These mice maintain physiologically sufficient insulin levels and do not develop diabetes or sustained hyperglycemia (Templeman et al. 2015). Female *Ins1*^-/-^ *;Ins2*^+/+^ and *Ins1^-/-^;Ins2^+/-^* littermates were weaned at 3–4 weeks of age and distributed between two dietary conditions, with weaned body weights approximately matched between diet groups. From weaning onward, mice were fed either a moderate-energy chow diet (CD; 10% fat Research Diets D12450K, New Brunswick, NJ, USA) or a high-energy, 45 kcal% fat, high-sucrose formulation (Research Diets D12451, New Brunswick, NJ, USA), both provided *ad libitum* except during designated fasting periods (Fig. 2A). Animals were housed under a 12:12 h light–dark cycle at 21°C.

### Fasting and glucose-stimulated insulin, glucose tolerance, and insulin sensitivity

Mice were fasted for 4 hours during the light period in clean cages to ensure a postprandial state. For intraperitoneal glucose tolerance tests (GTT), fasted mice received an intraperitoneal injection of sterile 20% glucose (2 g/kg body weight in 1X PBS) while for insulin tolerance tests (ITT), fasted mice were injected with 0.75 U/kg insulin analogue Humalog (Eli Lilly, Indianapolis, IN, USA). Blood glucose levels were measured at baseline (0 min) and at 15, 30-, 60-, 90-, and 120-minutes post-injection using a OneTouch Ultra2 or Ultra Mini glucometer. Fasting or glucose-stimulated insulin secretion (GSIS) was similarly tested after a 4-hour fast, at which time blood glucose levels were measured and a blood sample was collected. For glucose-stimulated insulin measurements, mice were given 2 g/kg glucose intraperitoneally, and blood samples were additionally collected at 15-, and 30-minutes post-injection. Plasma was isolated by centrifugation (13,000 rpm, 10 min, 4°C) and stored at –80°C until analysis. Insulin concentrations were quantified using a mouse ultrasensitive insulin ELISA kit (Alpco Diagnostics, Cat #80-INSMSU-E01). All assays were separated from one another by at least a week to allow mice to regain lost blood volume.

### Estrous cyclicity

Estrous cyclicity was monitored via daily vaginal smears collected between 08:00 am and 09:00 am over a 14-day period. A saline lavage (∼50 μL of sterile 1X PBS) was gently flushed 4–5 times over the vaginal opening. Vaginal cells were collected on microscope slides, and stained with 0.1% crystal violet (ThermoFisher Scientific, Cat #B12932.14). Slides were imaged using a Nikon Eclipse Ti2 microscope, and images were blinded and randomized prior to analysis. Estrous cycle stage was determined based on the relative abundance and morphology of vaginal cell types, including nucleated epithelial cells, cornified squamous epithelial cells, and leukocytes (Byers et al. 2012). Proestrus was characterized by nucleated epithelial cells, estrus by mostly cornified squamous epithelial cells, metestrus by a mixture of all three cell types, and diestrus by leukocytes and scattered nucleated epithelial cells.

### Hormone profiling

Plasma levels of Anti-Mullerian hormone (AMH) and follicle-stimulating (FSH) were measured in plasma collected in the morning on the first day of diestrus. Blood was collected from the saphenous vein and centrifuged at 13000 rpm for 10 minutes at 4°C to isolate plasma. AMH levels were quantified with an AMH ELISA (AL-113) from Ansh Labs and FSH concentrations were determined with an Abnova FSH ELISA kit (KA2278).

### Breeding performance

Female mice were paired individually with wild-type 2-month-old C57BL/6J males for periodic breeding performance trials after each set of metabolic and estrous cycling assessments. After being paired with male mice, the females were weighed every 3–4 days to monitor for pregnancy, which was signified by >10% increase in body weight along with visual signs of gestation. As parturition approached, breeding cages were checked every day for the presence of pups. Males remained with females for up to four weeks in the breeding performance trials if there was no pregnancy detected, or were removed earlier if pregnancy was confirmed and sustained for at least one week. If no pregnancy was detected within the four-week breeding trial, males were removed, and the females were returned to their original group housing. Litter sizes were determined from litters that have not include any cannibalization of pups.

### Ovarian histology

Ovaries were collected immediately after euthanasia via isoflurane inhalation and cervical dislocation, and were fixed in 4% paraformaldehyde at 4°C for 24 hours. Following fixation, ovaries were washed and stored in 70% ethanol at 4°C until processing. Fixed ovaries were then dehydrated through a graded ethanol series, embedded in paraffin, and sectioned at 5 µm thickness (Wax-it histology services, Vancouver, Canada). At least 10 sections were collected per mouse at equally distributed intervals spanning the ovary. Sections were stained with hematoxylin and eosin (H&E) or picrosirius red (PSR). Brightfield images were acquired using a Nikon Eclipse Ti2 microscope. Follicle subtypes were determined as previously described (Isola et al. 2020; Kim et al. 2025) in images that were blinded and randomized prior to assessment. Briefly, follicles were classified as primary if they were enclosed by one layer of cuboidal granulosa cells, and categorized as secondary if they were surrounded by more than one layer of cuboidal granulosa cells and had no clear antral cavity. Antral follicles were characterized by a layer of cumulus granulosa cells surrounding the oocyte and a visible antral space. Corpora lutea were identified by the presence of large clusters of granulosa lutein cells. Haemorrhagic corpora lutea were identified by the presence of red blood cells in its interstitial spaces, and we calculated the average percentage of corpora lutea that were haemorrhagic per mouse in ovary sections from 15 month-old mice. Fibrotic area was quantified by thresholding images of PSR-stained sections in ImageJ and calculating the precent of PSR-positive area relative to the rest of the ovary (Duncan and Pritchard 2024). To quantify multinucleated epithelial giant cells (MNGCs), we overlaid a grid over brightfield images of H&E-stained ovary sections that were blinded and randomized. For each section, we recorded the number of grids occupied by the ovary, and the number of those grids containing MNGCs. The grid size and position were unchanged between images. MNGCs were identified by their light brown foamy appearance as described in (Briley et al. 2016; Foley et al. 2021). Only mice with visible MNGCs in their ovary sections were included in this MNGC analysis.

All mice used for ovarian histology were from Facility A, either collected from an independently aged cohort (9 months), or collected after the last round of breeding trials (15 months).

### Statistical analyses

Statistical analyses were performed in GraphPad Prism 10 software and R studio (R version 4.1.1). For most variables in the mouse experiments, two-way ANOVAs were used to assess differences between groups based on the factors of diet and genotype. Detection of a statistically significant interaction between factors was followed by a one-way ANOVA with Bonferroni post-hoc corrections applied. Fisher’s exact tests were used to compare conception rates and pregnancy outcomes. To compare incidence of haemorrhagic corpora lutea between groups, we used the Kruksal-Wallis with Dunn’s post hoc analysis and Benjamani-Hochberg correction. To analyze MNGC counts, we used a generalized linear mixed model with diet, genotype, and their interaction as fixed effects, and animal_ID as a random intercept.

Participant data from the Study of Women’s Health Across the Nation were analysed through multiple linear regression using log-transformed insulin values and age at final menstrual period (FMP). FMP was estimated from the baseline age of the participant and the last self-reported menstrual cycle followed by a year of amenorrhea. To estimate adjusted mean FMP age across exposure strata, insulin levels were divided into quartiles (Q1–Q4) and unadjusted and covariate-adjusted models were fit with insulin as a predictor. Covariates included self-reported race/ethnicity, smoking status, and income at the baseline visit, as well as clinic-assessed BMI and fasting glucose levels.

The significance level was set at *p* ≤ 0.05 across all analyses.

## Acknowledgments

We thank Emma Houston and other members of the Templeman lab for valued discussions and experiment assistance throughout this study, and George Brownrigg and other members of the Johnson lab for additional assistance with mouse experiments at the University of British Columbia. This project was supported by funding to N.M.T. from the Canadian Institutes of Health Research (PJT - 183618) and the Banting Research Foundation (Discovery Award #2021-1427). N.M.T. is a Tier 2 Canada Research Chair in Metabolic Determinants of Reproduction and Aging, and a Michael Smith Health Research BC Scholar. L.G.H. was supported by the Canadian Islet Research and Training Network and Michael Smith Health Research BC.

We used publicly available data from the Study of Women’s Health Across the Nation (SWAN), which has had grant support through the National Institutes of Health (NIH), Department of Health and Human Services (DHHS), the National Institute on Aging (NIA), the National Institute of Nursing Research (NINR), and the NIH Office of Research on Women’s Health (ORWH). We thank all women who participated in SWAN, as well as the study staff at each site and the primary investigators and administrators of the SWAN study. We also appreciated guidance and discussions with Sarah Gregory, Graciela Muniz-Terrera, and Andrea Piccinin while conducting our secondary analyses of SWAN data.

## Author contributions

N.M.T. conceptualized the study, and N.M.T. and F.A. designed experiments. F.A. conducted secondary analyses of SWAN data. J.D.J. provided mouse models and reagents, as well as input on experimental design. F.A., L.H., and X.H. performed mouse experiments, and F.A. analyzed and interpreted data together with N.M.T. The manuscript was written by F.A. and N.M.T., and all authors provided editorial feedback and approved the final version of the paper.

## Declaration of interests

The authors declare no competing interests.

## Supplemental information

**Table S1:** Demographics and characteristics of analytic sample from the Study of Women’s Health Across the Nation (SWAN). Unless otherwise indicated, data are presented as count (percentages) for categorical variables, and as mean (SD) for continuous variables.

|  |  |
| --- | --- |
| **Analytic sample size** (of total SWAN enrollment) | 646 (of 3302 SWAN participants total) |
| **Self-reported race and ethnicity** |  |
| Caucasian/White Non-Hispanic | 352 (54.5%) |
| Black/African American | 126 (19.5%) |
| Japanese/Japanese American | 89 (13.8%) |
| Chinese/Chinese American | 79 (12.2%) |
| **Income bracket** |  |
| < \$20,000 | 38 (5.9%) |
| \$20,000-49,000 | 211 (32.7%) |
| \$50,000-99,000 | 278 (43.0%) |
| > \$100,000 | 119 (18.4%) |
| **Smoking status** | 239 (37%) current smokers |
| **Fasting insulin at 46-47 years old (μIU/mL)** | 9.94 (SD 6.34) |
| **BMI at 46-47 years old (kg/m<sup>2</sup>)** | 26.8 (SD 6.6) |
| **Fasting glucose at 46-47 years old (mg/dL)** | 89.6 (SD 9.8) |
| **Age at final menstrual period** | 51.2 (SD 2.1) |

**Table S2:** Associations between age-47 fasting insulin levels and age of menopause. Associations are results of linear regression models between log-transformed fasting insulin values at 46 or 47 years old and age of final menstrual period. Unadjusted model: simple linear regression; model adjusted for demographics: multiple linear regression adjusted for self-reported race/ethnicity, income, and smoking status; model adjusted for demographics and metabolic confounders: multiple linear regression adjusted for demographic covariates (self-reported race/ethnicity, income, and smoking status) as well as metabolic confounders of clinic-assessed BMI and fasting glucose levels, for both insulin levels and age of final menstrual period. Ordinary least squares estimation was applied. Bold text signifies p ≤ 0.05.

|  |  |
| --- | --- |
| **Unadjusted model** |  |
| $\beta$ coefficient | -0.177 |
| p-value | 0.292 |
| Adjusted R<sup>2</sup> value | 0.0002 |
| **Model adjusted for demographic covariates:**<br>**self-reported race/ethnicity + income + smoking status** |  |
| $\beta$ coefficient | -0.127 |
| p-value | 0.466 |
| Adjusted R<sup>2</sup> value | 0.0142 |
| **Model adjusted for demographic covariates**<br>**(self-reported race/ethnicity + income + smoking status)**<br>**and for metabolic confounders:**<br>**BMI and fasting glucose at age 46-47** |  |
| $\beta$ coefficient | **-0.471** |
| p-value | **0.033** |
| Adjusted R<sup>2</sup> value | **0.0229** |

**Fig S1.**
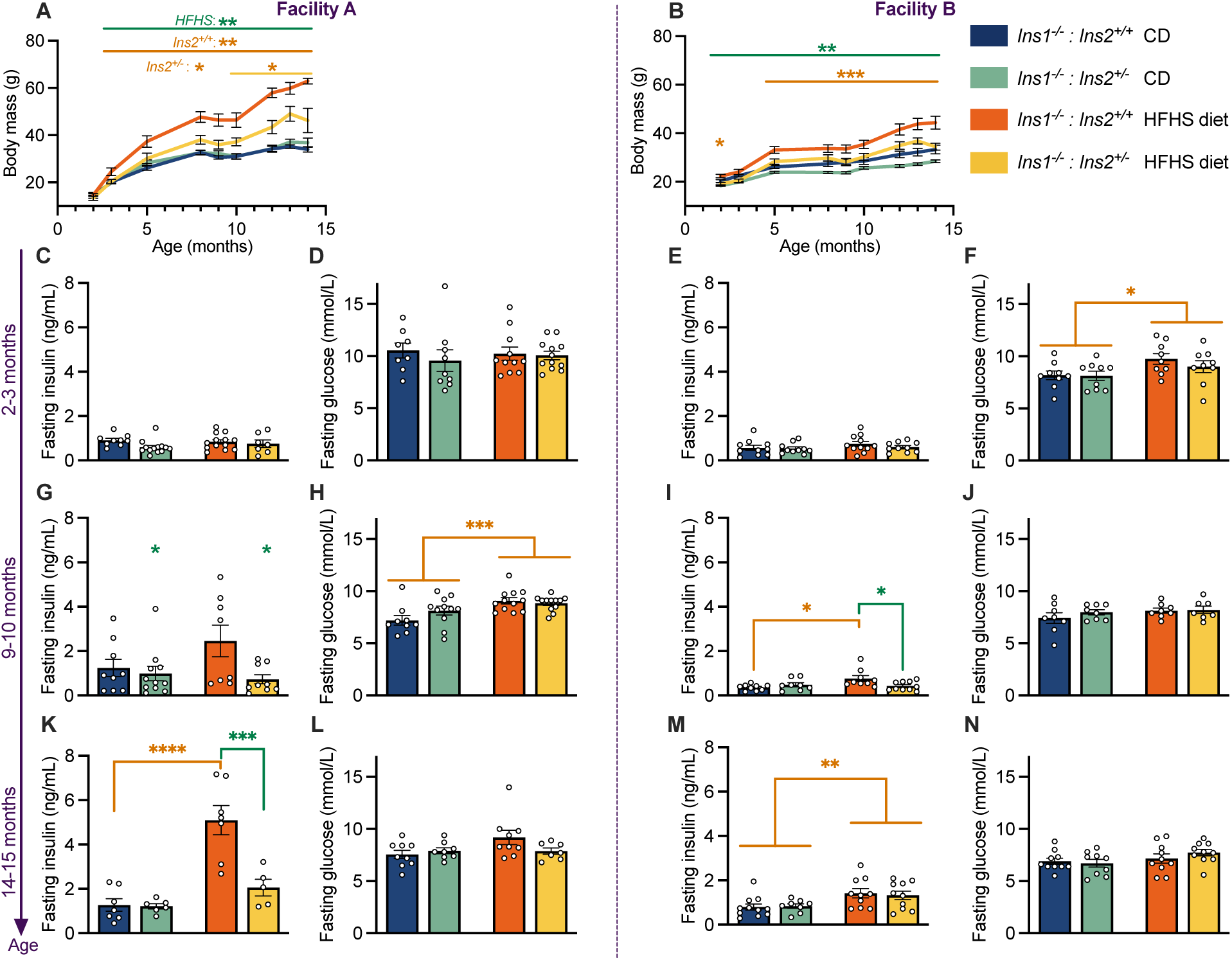
Similar trends in metabolic parameters across both facilities. Longitudinal body weight measurements for (A) Facility A mouse cohort and (B) Facility B mouse cohort. (C, G, K) Fasting plasma insulin levels at 2-3, 9-10, and 14-15 months of age for Facility A. (D, H, L) Fasting blood glucose concentrations at 2-3, 9-10, and 14-15 months of age for Facility A. (E, I, M) Fasting plasma insulin levels at 2-3, 9-10, and 14-15 months of age for Facility B. (F, J, N) Fasting blood glucose concentrations at 2-3, 9-10, and 14-15 months of age for Facility B. Data are presented as means and SEM; scatter points represent individual mice. Statistical analyses were performed using two-way ANOVA to evaluate the effects of genotype and diet (performed at each age point for body mass data). When significant interactions were detected, one-way ANOVAs with Bonferroni post hoc corrections were conducted. Orange asterisks indicate a statistically significant effect of diet, within one of the genotypes (in case of a statistical interaction) where indicated; green asterisks indicate a statistically significant genotype effect, within one of the diets (in case of a statistical interaction) where indicated. * *p* < 0.05; ** *p* < 0.01; *** *p* < 0.001; **** *p* < 0.0001. n = 4-11 from Facility A; n = 8-10 from Facility B.

## Notes

### Competing Interest Statement

The authors have declared no competing interest.

## References

Acevedo, Nicole, Jun Ding, and Gary D. Smith. 2007. “Insulin Signaling in Mouse Oocytes.” Biology of Reproduction 77 (5): 872–79. 10.1095/biolreprod.107.060152.

Athar, Faria, Muskan Karmani, and Nicole M. Templeman. 2024. “Metabolic Hormones Are Integral Regulators of Female Reproductive Health and Function.” Bioscience Reports 44 (1): BSR20231916. 10.1042/BSR20231916.

Bajalan, Zahra, Zainab Alimoradi, and Farnoosh Moafi. 2019. “Nutrition as a Potential Factor of Primary Dysmenorrhea: A Systematic Review of Observational Studies.” Gynecologic and Obstetric Investigation 84 (3): 209–24. 10.1159/000495408.

Balough, Julia L., Shweta S. Dipali, Karen Velez, T. Rajendra Kumar, and Francesca E. Duncan. 2024. “Hallmarks of Female Reproductive Aging in Physiologic Aging Mice.” Nature Aging 4 (12): 1711–30. 10.1038/s43587-024-00769-y.

Bird, Stephen R, and John A Hawley. 2017. “Update on the Effects of Physical Activity on Insulin Sensitivity in Humans.” BMJ Open Sport — Exercise Medicine 2 (1): e000143. 10.1136/bmjsem-2016-000143.

Blüher, Matthias, M. Dodson Michael, Odile D. Peroni, et al. 2002. “Adipose Tissue Selective Insulin Receptor Knockout Protects against Obesity and Obesity-Related Glucose Intolerance.” Developmental Cell 3 (1): 25–38. 10.1016/s1534-5807(02)00199-5.

Briley, Shawn M., Susmita Jasti, Jennifer M. McCracken, et al. 2016. Reproductive Age-Associated Fibrosis in the Stroma of the Mammalian Ovary. Reproduction. September 1. 10.1530/REP-16-0129.

Brothers, Kathryn J., Sheng Wu, Sara A. Divall, et al. 2010. “Rescue of Obesity-Induced Infertility in Female Mice Due to a Pituitary-Specific Knockout of the Insulin Receptor.” Cell Metabolism 12 (3): 295–305. 10.1016/j.cmet.2010.06.010.

Brown, Elise C., Barry A. Franklin, Judith G. Regensteiner, and Kerry J. Stewart. 2020. “Effects of Single Bout Resistance Exercise on Glucose Levels, Insulin Action, and Cardiovascular Risk in Type 2 Diabetes: A Narrative Review.” Journal of Diabetes and Its Complications 34 (8): 107610. 10.1016/j.jdiacomp.2020.107610.

Brüning, J. C., D. Gautam, D. J. Burks, et al. 2000. “Role of Brain Insulin Receptor in Control of Body Weight and Reproduction.” Science (New York, N.Y.) 289 (5487): 2122–25. 10.1126/science.289.5487.2122.

Byers, Shannon L., Michael V. Wiles, Sadie L. Dunn, and Robert A. Taft. 2012. “Mouse Estrous Cycle Identification Tool and Images.” PLoS ONE 7 (4): e35538. 10.1371/journal.pone.0035538.

Carnethon, Mercedes R., Latha P. Palaniappan, Cecil M. Burchfiel, Frederick L. Brancati, and Stephen P. Fortmann. 2002. “Serum Insulin, Obesity, and the Incidence of Type 2 Diabetes in Black and White Adults: The Atherosclerosis Risk in Communities Study: 1987-1998.” Diabetes Care 25 (8): 1358–64. 10.2337/diacare.25.8.1358.

Castrillon, Diego H., Lili Miao, Ramya Kollipara, James W. Horner, and Ronald A. DePinho. 2003. “Suppression of Ovarian Follicle Activation in Mice by the Transcription Factor Foxo3a.” Science 301 (5630): 215–18. 10.1126/science.1086336.

Chavarro, Jorge E., Janet W. Rich-Edwards, Bernard A. Rosner, and Walter C. Willett. 2007. “Diet and Lifestyle in the Prevention of Ovulatory Disorder Infertility.” Obstetrics and Gynecology 110 (5): 1050–58. 10.1097/01.AOG.0000287293.25465.e1.

Cheng, Jinmei, Yinchuan Li, Yan Zhang, Xiuxia Wang, Fei Sun, and Yixun Liu. 2020. “Conditional Deletion of *Wntless* in Granulosa Cells Causes Impaired Corpora Lutea Formation and Subfertility.” Aging 13 (1): 1001–16. 10.18632/aging.202222.

Collins, Gretchen, Bansari Patel, Suruchi Thakore, and James Liu. 2017. “Primary Ovarian Insufficiency: Current Concepts.” Southern Medical Journal 110 (3): 147–53. 10.14423/SMJ.0000000000000611.

Converse, Aubrey, Madeline J. Perry, Shweta S. Dipali, et al. 2025. “Multinucleated Giant Cells Are Hallmarks of Ovarian Aging with Unique Immune and Degradation-Associated Molecular Signatures.” PLOS Biology 23 (6): e3003204. 10.1371/journal.pbio.3003204.

Cust, Anne E., Naomi E. Allen, Sabina Rinaldi, et al. 2007. “Serum Levels of C-Peptide, IGFBP-1 and IGFBP-2 and Endometrial Cancer Risk; Results from the European Prospective Investigation into Cancer and Nutrition.” International Journal of Cancer 120 (12): 2656–64. 10.1002/ijc.22578.

Davis, Susan R., JoAnn Pinkerton, Nanette Santoro, and Tommaso Simoncini. 2023. “Menopause—Biology, Consequences, Supportive Care, and Therapeutic Options.” *Cell*, ahead of print, September 6. 10.1016/j.cell.2023.08.016.

Di Berardino, Chiara, Urte Barceviciute, Chiara Camerano Spelta Rapini, et al. 2024. “High-Fat Diet-Negative Impact on Female Fertility: From Mechanisms to Protective Actions of Antioxidant Matrices.” Frontiers in Nutrition 11 (June). 10.3389/fnut.2024.1415455.

Diamanti-Kandarakis, Evanthia, and Andrea Dunaif. 2012. “Insulin Resistance and the Polycystic Ovary Syndrome Revisited: An Update on Mechanisms and Implications.” Endocrine Reviews 33 (6): 981–1030. 10.1210/er.2011-1034.

DiVall, Sara A., Danny Herrera, Bonnie Sklar, et al. 2015. “Insulin Receptor Signaling in the GnRH Neuron Plays a Role in the Abnormal GnRH Pulsatility of Obese Female Mice.” PLOS ONE 10 (3): e0119995. 10.1371/journal.pone.0119995.

Duffy, Diane M., CheMyong Ko, Misung Jo, Mats Brannstrom, and Thomas E. Curry. 2019. “Ovulation: Parallels With Inflammatory Processes.” Endocrine Reviews 40 (2): 369–416. 10.1210/er.2018-00075.

Duncan, Francesca E., and Michele T. Pritchard. 2024. Picrosirius Red (PSR) Staining and Quantification in Mouse Ovaries. November 9. https://www.protocols.io/view/picrosirius-red-psr-staining-and-quantification-in-dqwb5xan.

Dunneram, Yashvee, Darren Charles Greenwood, Victoria J. Burley, and Janet E. Cade. 2018. “Dietary Intake and Age at Natural Menopause: Results from the UK Women’s Cohort Study.” Journal of Epidemiology and Community Health 72 (8): 733–40. 10.1136/jech-2017-209887.

Duvillié, Bertrand, Nathalie Cordonnier, Louise Deltour, et al. 1997. “Phenotypic Alterations in Insulin-Deficient Mutant Mice.” Proceedings of the National Academy of Sciences of the United States of America 94 (10): 5137–40. 10.1073/pnas.94.10.5137.

Faddy, M. J., and R. G. Gosden. 1996. “A Model Conforming the Decline in Follicle Numbers to the Age of Menopause in Women.” Human Reproduction 11 (7): 1484–86. 10.1093/oxfordjournals.humrep.a019422.

Foley, K. Grace, Michele T. Pritchard, and Francesca E. Duncan. 2021. “Macrophage Multinucleated Giant Cells – Hallmarks of the Aging Ovary.” Reproduction (Cambridge, England) 161 (2): V5–9. 10.1530/REP-20-0489.

Garcia, Driele N., Tatiana D. Saccon, Jorgea Pradiee, et al. 2019. “Effect of Caloric Restriction and Rapamycin on Ovarian Aging in Mice.” GeroScience 41 (4): 395–408. 10.1007/s11357-019-00087-x.

Garruti, Gabriella, Raffaella Depalo, and Maria De Angelis. 2019. “Weighing the Impact of Diet and Lifestyle on Female Reproductive Function.” Current Medicinal Chemistry 26 (19): 3584–92. 10.2174/0929867324666170518101008.

Gaskins, Audrey J., Feiby L. Nassan, Yu-Han Chiu, et al. 2019. “Dietary Patterns and Outcomes of Assisted Reproduction.” American Journal of Obstetrics and Gynecology 220 (6): 567.e1-567.e18. 10.1016/j.ajog.2019.02.004.

Ghasemi, Asghar, Maryam Tohidi, Arash Derakhshan, Mitra Hasheminia, Fereidoun Azizi, and Farzad Hadaegh. 2015. “Cut-off Points of Homeostasis Model Assessment of Insulin Resistance, Beta-Cell Function, and Fasting Serum Insulin to Identify Future Type 2 Diabetes: Tehran Lipid and Glucose Study.” Acta Diabetologica 52 (5): 905–15. 10.1007/s00592-015-0730-3.

Gold, Ellen B., Sybil L. Crawford, Nancy E. Avis, et al. 2013. “Factors Related to Age at Natural Menopause: Longitudinal Analyses from SWAN.” American Journal of Epidemiology 178 (1): 70–83. 10.1093/aje/kws421.

Grieger, Jessica A., Luke E. Grzeskowiak, Tina Bianco-Miotto, et al. 2018. “Pre-Pregnancy Fast Food and Fruit Intake Is Associated with Time to Pregnancy.” *Human Reproduction (Oxford*, England) 33 (6): 1063–70. 10.1093/humrep/dey079.

Gu, Mengqing, Yibo Wang, and Yang Yu. 2024. “Ovarian Fibrosis: Molecular Mechanisms and Potential Therapeutic Targets.” Journal of Ovarian Research 17 (1): 139. 10.1186/s13048-024-01448-7.

Guan, Yu, Qian Liu, Zhimin Deng, et al. 2025. “Association of Social Determinants of Health and Age at Menopause: NHANES 1999–2018 Observational Study.” Human Reproduction Open 2025 (3): hoaf050. 10.1093/hropen/hoaf050.

Gunter, Marc J., Donald R. Hoover, Herbert Yu, et al. 2008. “A Prospective Evaluation of Insulin and Insulin-like Growth Factor-I as Risk Factors for Endometrial Cancer.” *Cancer Epidemiology, Biomarkers & Prevention: A Publication of the American Association for Cancer Research*, Cosponsored by the American Society of Preventive Oncology 17 (4): 921–29. 10.1158/1055-9965.EPI-07-2686.

Hall, Janet E. 2015. “Endocrinology of the Menopause.” Endocrinology and Metabolism Clinics of North America 44 (3): 485–96. 10.1016/j.ecl.2015.05.010.

Hohos, Natalie M., Kirstin J. Cho, Delaney C. Swindle, and Malgorzata E. Skaznik-Wikiel. 2018. “High-Fat Diet Exposure, Regardless of Induction of Obesity, Is Associated with Altered Expression of Genes Critical to Normal Ovulatory Function.” Molecular and Cellular Endocrinology 470 (July): 199–207. 10.1016/j.mce.2017.10.016.

Hohos, Natalie M, and Malgorzata E Skaznik-Wikiel. 2017. “High-Fat Diet and Female Fertility.” Endocrinology 158 (8): 2407–19. 10.1210/en.2017-00371.

Hong, Jae Seok, Sang Wook Yi, Hee Chung Kang, et al. 2007. “Age at Menopause and Cause-Specific Mortality in South Korean Women: Kangwha Cohort Study.” Maturitas 56 (4): 411–19. 10.1016/j.maturitas.2006.11.004.

Igarashi, Hideki, Toshifumi Takahashi, and Satoru Nagase. 2015. “Oocyte Aging Underlies Female Reproductive Aging: Biological Mechanisms and Therapeutic Strategies.” Reproductive Medicine and Biology 14 (4): 159–69. 10.1007/s12522-015-0209-5.

Isola, José V. V., Bianka M. Zanini, Silvana Sidhom, et al. 2020. “17α-Estradiol Promotes Ovarian Aging in Growth Hormone Receptor Knockout Mice, but Not Wild-Type Littermates.” Experimental Gerontology 129 (January): 110769. 10.1016/j.exger.2019.110769.

Johnson, Tammie M., and James R. Churilla. 2025. “Trends in Mean Serum Insulin and Hyperinsulinemia among US Adults without Diabetes 1999–2018.” Journal of Diabetes and Its Complications 39 (11): 109159. 10.1016/j.jdiacomp.2025.109159.

Karayiannis, Dimitrios, Meropi D. Kontogianni, Christina Mendorou, Minas Mastrominas, and Nikos Yiannakouris. 2018. “Adherence to the Mediterranean Diet and IVF Success Rate among Non-Obese Women Attempting Fertility.” *Human Reproduction (Oxford*, England) 33 (3): 494–502. 10.1093/humrep/dey003.

Khalil, N., K. Sutton-Tyrrell, E. S. Strotmeyer, et al. 2011. “Menopausal Bone Changes and Incident Fractures in Diabetic Women: A Cohort Study.” Osteoporosis International : A Journal Established as Result of Cooperation between the European Foundation for Osteoporosis and the National Osteoporosis Foundation of the USA 22 (5): 1367–76. 10.1007/s00198-010-1357-4.

Kim, Minhoo, Rajyk Bhala, Justin Wang, et al. 2025. “Systematic Characterization of the Ovarian Landscape across Mouse Menopause Models.” Preprint, bioRxiv, June 15. 10.1101/2025.06.10.658920.

Kim, YoonMyung, and HaNui Park. 2013. “Does Regular Exercise without Weight Loss Reduce Insulin Resistance in Children and Adolescents?” International Journal of Endocrinology 2013: 402592. 10.1155/2013/402592.

Kirwan, John P., Thomas P. J. Solomon, Daniel M. Wojta, Myrlene A. Staten, and John O. Holloszy. 2009. “Effects of 7 Days of Exercise Training on Insulin Sensitivity and Responsiveness in Type 2 Diabetes Mellitus.” American Journal of Physiology-Endocrinology and Metabolism 297 (1): E151–56. 10.1152/ajpendo.00210.2009.

Kitamura, Tadahiro, C. Ronald Kahn, and Domenico Accili. 2003. “Insulin Receptor Knockout Mice.” Annual Review of Physiology 65: 313–32. 10.1146/annurev.physiol.65.092101.142540.

Leroux, Loïc, Pierrette Desbois, Luciane Lamotte, et al. 2001. “Compensatory Responses in Mice Carrying a Null Mutation for Ins1 or Ins2.” Diabetes 50 (SUPPL. 1): 150–53. 10.2337/diabetes.50.2007.s150.

Li, Chaoyang, Earl S. Ford, Lisa C. McGuire, Ali H. Mokdad, Randie R. Little, and Gerald M. Reaven. 2006. “Trends in Hyperinsulinemia Among Nondiabetic Adults in the U.S.” Diabetes Care 29 (11): 2396–402. 10.2337/dc06-0289.

Luo, Shijing, Gunnar A. Kleemann, Jasmine M. Ashraf, Wendy M. Shaw, and Coleen T. Murphy. 2010. “TGF-β and Insulin Signaling Regulate Reproductive Aging via Oocyte and Germline Quality Maintenance.” Cell 143 (2): 299–312. 10.1016/j.cell.2010.09.013.

Matthews, Karen A., Sybil L. Crawford, Claudia U. Chae, et al. 2009. “Are Changes in Cardiovascular Disease Risk Factors in Midlife Women Due to Chronological Aging or to the Menopausal Transition?” Journal of the American College of Cardiology 54 (25): 2366–73. 10.1016/j.jacc.2009.10.009.

Mehra, Vrati M., Christy Costanian, Hugh McCague, Michael C. Riddell, and Hala Tamim. 2023. “The Association between Diabetes Type, Age of Onset, and Age at Natural Menopause: A Retrospective Cohort Study Using the Canadian Longitudinal Study on Aging.” Menopause 30 (1): 37. 10.1097/GME.0000000000002085.

Mehran, Arya E., Nicole M. Templeman, G. Stefano Brigidi, et al. 2012. “Hyperinsulinemia Drives Diet-Induced Obesity Independently of Brain Insulin Production.” Cell Metabolism 16 (6): 723–37. 10.1016/j.cmet.2012.10.019.

Mishina, Tappei, Namine Tabata, Tetsutaro Hayashi, et al. 2021. “Single-Oocyte Transcriptome Analysis Reveals Aging-Associated Effects Influenced by Life Stage and Calorie Restriction.” Aging Cell 20 (8): 1–11. 10.1111/acel.13428.

Mitro, Susanna D, L Elaine Waetjen, Catherine Lee, et al. 2024. “Diabetes and Uterine Fibroid Diagnosis in Midlife: Study of Women’s Health Across the Nation (SWAN).” The Journal of Clinical Endocrinology and Metabolism 110 (6): e1934–42. 10.1210/clinem/dgae625.

Mittendorfer, Bettina, James D. Johnson, Giovanni Solinas, and Per-Anders Jansson. 2024. “Insulin Hypersecretion as Promoter of Body Fat Gain and Hyperglycemia.” Diabetes 73 (6): 837–43. 10.2337/dbi23-0035.

Nagel, Gabriele, Hans Peter Altenburg, Alexandra Nieters, Paolo Boffetta, and Jakob Linseisen. 2005. “Reproductive and Dietary Determinants of the Age at Menopause in EPIC-Heidelberg.” Maturitas 52 (3–4): 337–47. 10.1016/j.maturitas.2005.05.013.

Nakae, J., Y. Kido, and D. Accili. 2001. “Distinct and Overlapping Functions of Insulin and IGF-I Receptors.” Endocrine Reviews 22 (6): 818–35. 10.1210/edrv.22.6.0452.

Odeleye, O. E., M. de Courten, D. J. Pettitt, and E. Ravussin. 1997. “Fasting Hyperinsulinemia Is a Predictor of Increased Body Weight Gain and Obesity in Pima Indian Children.” Diabetes.. 46 (8): 1341–45. 10.2337/diab.46.8.1341.

Ossewaarde, Marlies E., Michiel L. Bots, André L.M. Verbeek, et al. 2005. “Age at Menopause, Cause-Specific Mortality and Total Life Expectancy.” Epidemiology 16 (4): 556–62. 10.1097/01.ede.0000165392.35273.d4.

Ou, Xiang-Hong, Sen Li, Zhen-Bo Wang, et al. 2012. “Maternal Insulin Resistance Causes Oxidative Stress and Mitochondrial Dysfunction in Mouse Oocytes.” Human Reproduction 27 (7): 2130–45. 10.1093/humrep/des137.

Parkening, T. A., T. J. Collins, and E. R. Smith. 1982. “Plasma and Pituitary Concentrations of LH, FSH, and Prolactin in Aging C57BL/6 Mice at Various Times of the Estrous Cycle.” Neurobiology of Aging 3 (1): 31–35. 10.1016/0197-4580(82)90058-6.

Poretsky, L., N. A. Cataldo, Z. Rosenwaks, and L. C. Giudice. 1999. “The Insulin-Related Ovarian Regulatory System in Health and Disease.” Endocrine Reviews 20 (4): 535–82. 10.1210/edrv.20.4.0374.

Qiu, Xiaoliang, Hoangha Dao, Mengjie Wang, et al. 2015. “Insulin and Leptin Signaling Interact in the Mouse Kiss1 Neuron during the Peripubertal Period.” PloS One 10 (5): e0121974. 10.1371/journal.pone.0121974.

Qiu, Xiaoliang, Abigail R. Dowling, Joseph S. Marino, et al. 2013. “Delayed Puberty but Normal Fertility in Mice with Selective Deletion of Insulin Receptors from Kiss1 Cells.” Endocrinology 154 (3): 1337–48. 10.1210/en.2012-2056.

Ritenbaugh, Cheryl, Nicolette I Teufel-Shone, Mikel G Aickin, et al. 2003. “A Lifestyle Intervention Improves Plasma Insulin Levels among Native American High School Youth.” Preventive Medicine 36 (3): 309–19. 10.1016/S0091-7435(02)00015-4.

Roa-Díaz, Zayne Milena, Faina Wehrli, Irene Lambrinoudaki, et al. 2023. “Early Menopause and Cardiovascular Risk Factors: A Cross-Sectional and Longitudinal Study.” Menopause 30 (6): 599. 10.1097/GME.0000000000002184.

Savukoski, Susanna M., Eila T. J. Suvanto, Juha P. Auvinen, et al. 2021. “Onset of the Climacteric Phase by the Mid-Forties Associated with Impaired Insulin Sensitivity: A Birth Cohort Study.” Menopause 28 (1): 70. 10.1097/GME.0000000000001658.

Schoenaker, Danielle A. J. M., Caroline A. Jackson, Jemma V. Rowlands, and Gita D. Mishra. 2014. “Socioeconomic Position, Lifestyle Factors and Age at Natural Menopause: A Systematic Review and Meta-Analyses of Studies across Six Continents.” International Journal of Epidemiology 43 (5): 1542–62. 10.1093/ije/dyu094.

Sekhar, T.V.D. Sasi, Soumya Medarametla, Arifa Rahman, and Satya Sahi Adapa. 2015. “Early Menopause in Type 2 Diabetes – A Study from a South Indian Tertiary Care Centre.” Journal of Clinical and Diagnostic Research : JCDR 9 (10): OC08–OC10. 10.7860/JCDR/2015/14181.6628.

Selesniemi, Kaisa, Ho Joon Lee, and Jonathan L. Tilly. 2008. “Moderate Caloric Restriction Initiated in Rodents during Adulthood Sustains Function of the Female Reproductive Axis into Advanced Chronological Age.” Aging Cell 7 (5): 622–29. 10.1111/j.1474-9726.2008.00409.x.

Shadyab, Aladdin H., Caroline A. MacEra, Richard A. Shaffer, et al. 2017. “Ages at Menarche and Menopause and Reproductive Lifespan as Predictors of Exceptional Longevity in Women: The Women’s Health Initiative.” Menopause 24 (1): 35–44. 10.1097/GME.0000000000000710.

Skaznik-Wikiel, Malgorzata E., Delaney C. Swindle, Amanda A. Allshouse, Alex J. Polotsky, and James L. McManaman. 2016. “High-Fat Diet Causes Subfertility and Compromised Ovarian Function Independent of Obesity in Mice.” Biology of Reproduction 94 (5): 108. 10.1095/biolreprod.115.137414.

Sowers, Maryfran, Sybil L. Crawford, Barbara Sternfeld, et al. 2000. “CHAPTER 11 - SWAN: A Multicenter, Multiethnic, Community-Based Cohort Study of Women and the Menopausal Transition.” In Menopause, edited by ROGERIO A. Lobo, JENNIFER Kelsey, and ROBERT Marcus. Academic Press. 10.1016/B978-012453790-3/50012-3.

Speakman, John R. 2019. “Use of High-Fat Diets to Study Rodent Obesity as a Model of Human Obesity.” International Journal of Obesity 43 (8): 1491–92. 10.1038/s41366-019-0363-7.

Sun, Wanwan, Jieli Lu, Shengli Wu, et al. 2016. “Association of Insulin Resistance with Breast, Ovarian, Endometrial and Cervical Cancers in Non-Diabetic Women.” American Journal of Cancer Research 6 (10): 2334–44.

Templeman, Nicole M., Susanne M. Clee, and James D. Johnson. 2015. “Suppression of Hyperinsulinaemia in Growing Female Mice Provides Long-Term Protection against Obesity.” Diabetologia 58 (10): 2392–402. 10.1007/s00125-015-3676-7.

Templeman, Nicole M., Stephane Flibotte, Jenny H.L. Chik, et al. 2017. “Reduced Circulating Insulin Enhances Insulin Sensitivity in Old Mice and Extends Lifespan.” Cell Reports 20 (2): 451–63. 10.1016/j.celrep.2017.06.048.

Templeman, Nicole M., Søs Skovsø, Melissa M. Page, Gareth E. Lim, and James D. Johnson. 2017. “A Causal Role for Hyperinsulinemia in Obesity.” Journal of Endocrinology 232 (3): R173–83. 10.1530/JOE-16-0449.

Thomas, Dylan D, Barbara E Corkey, Nawfal W Istfan, and Caroline M Apovian. 2019. “Hyperinsulinemia: An Early Indicator of Metabolic Dysfunction.” Journal of the Endocrine Society 3 (9): 1727–47. 10.1210/js.2019-00065.

Tissenbaum, Heidi A., and Gary Ruvkun. 1998. “An Insulin-like Signaling Pathway Affects Both Longevity and Reproduction in Caenorhabditis Elegans.” Genetics 148 (2): 703–17. 10.1093/genetics/148.2.703.

Tricò, Domenico, Andrea Natali, Silva Arslanian, Andrea Mari, and Ele Ferrannini. 2018. “Identification, Pathophysiology, and Clinical Implications of Primary Insulin Hypersecretion in Nondiabetic Adults and Adolescents.” JCI Insight 3 (24): e124912. 10.1172/jci.insight.124912.

Turvey, Samuel J., Martin J. McPhillie, Mark T. Kearney, Stephen P. Muench, Katie J. Simmons, and Colin W. G. Fishwick. 2022. “Recent Developments in the Structural Characterisation of the IR and IGF1R: Implications for the Design of IR–IGF1R Hybrid Receptor Modulators.” RSC Medicinal Chemistry 13 (4): 360–74. 10.1039/d1md00300c.

Umehara, Takashi, Yasmyn E. Winstanley, Eryk Andreas, et al. 2022. “Female Reproductive Life Span Is Extended by Targeted Removal of Fibrotic Collagen from the Mouse Ovary.” Science Advances 8 (24): eabn4564. 10.1126/sciadv.abn4564.

Vella, Veronica, Agostino Milluzzo, Nunzio Massimo Scalisi, Paolo Vigneri, and Laura Sciacca. 2018. “Insulin Receptor Isoforms in Cancer.” International Journal of Molecular Sciences 19 (11): 3615. 10.3390/ijms19113615.

Vliet, Stephan van, Han-Chow E. Koh, Bruce W. Patterson, et al. 2020. “Obesity Is Associated With Increased Basal and Postprandial β-Cell Insulin Secretion Even in the Absence of Insulin Resistance.” Diabetes 69 (10): 2112–19. 10.2337/db20-0377.

Wellons, Melissa F., Juliana J. Matthews, and Catherine Kim. 2017. “Ovarian Aging in Women with Diabetes: An Overview.” Maturitas 96 (February): 109–13. 10.1016/j.maturitas.2016.11.019.

Willis, Sydney K., Lauren A. Wise, Amelia K. Wesselink, et al. 2020. “Glycemic Load, Dietary Fiber, and Added Sugar and Fecundability in 2 Preconception Cohorts.” The American Journal of Clinical Nutrition 112 (1): 27–38. 10.1093/ajcn/nqz312.

Wu, Sheng, Sara Divall, Amanda Nwaopara, et al. 2014. “Obesity-Induced Infertility and Hyperandrogenism Are Corrected by Deletion of the Insulin Receptor in the Ovarian Theca Cell.” Diabetes 63 (4): 1270–82. 10.2337/db13-1514.

Zaniker, Emily J., Elnur Babayev, and Francesca E. Duncan. 2023. “Common Mechanisms of Physiological and Pathological Rupture Events in Biology: Novel Insights into Mammalian Ovulation and Beyond.” Biological Reviews 98 (5): 1648–67. 10.1111/brv.12970.

Zhang, Qiao-Li, Yan Wang, Jian-Sheng Liu, and Yan-Zhi Du. 2022. “Effects of Hypercaloric Diet-Induced Hyperinsulinemia and Hyperlipidemia on the Ovarian Follicular Development in Mice.” The Journal of Reproduction and Development 68 (3): 173–80. 10.1262/jrd.2021-132.

